# Modelling the within-host spread of SARS-CoV-2 infection, and the subsequent immune response, using a hybrid, multiscale, individual-based model. *Part I: Macrophages*

**DOI:** 10.1101/2022.05.06.490883

**Authors:** C. F. Rowlatt, M. A. J. Chaplain, D. J. Hughes, S. H. Gillespie, D. H. Dockrell, I. Johannessen, R. Bowness

## Abstract

Individual responses to SARS-CoV-2 infection vary significantly, ranging from mild courses of infection that do not require hospitalisation to the development of disease which not only requires hospitalisation but can be fatal. Whilst many immunological studies have revealed fundamental insights into SARS-CoV-2 infection and COVID-19, mathematical and computational modelling can offer an additional perspective and enhance understanding. The majority of mathematical models for the within-host spread of SARS-CoV-2 infection are ordinary differential equations, which neglect spatial variation. In this article, we present a hybrid, multiscale, individual-based model to study the within-host spread of SARS-CoV-2 infection. The model incorporates epithelial cells (each containing a dynamical model for viral entry and replication), macrophages and a subset of cytokines. We investigate the role of increasing initial viral deposition, increasing delay in type I interferon secretion from epithelial cells (as well as the magnitude of secretion), increasing macrophage virus internalisation rate and macrophage activation, on the spread of infection.

## 1 Introduction

Individual responses to severe acute respiratory syndrome coronavirus 2 (SARS-CoV-2) infection, the causative agent of the COVID-19 pandemic, are heterogeneous; for the original (Wuhan) strain, and subsequent Alpha to Delta variants, the majority of cases present a mild course of infection (with a significant proportion being asymptomatic) and do not require hospitalisation, whilst some infections develop into coronavirus disease (COVID-19), which not only require hospitalisation but can be fatal.

A dysregulated immune response is associated with severe COVID-19, characterised by an aberrant immune cell distribution, as well as elevated inflammatory profiles, [1, 2, 3, 4, 5]. Indeed, early studies demonstrated a loss of resident alveolar macrophages, an increase in inflammatory monocyte-derived macrophages and neutrophils, loss of regulatory T cell responses, and an overall decrease in lymphocytes, in patients with severe disease, as well as an increase in proinflammatory cytokines (such as interleukin 1 (IL-1), interleukin 6 (IL-6) and tumor necrosis factor (TNF-*α*)) [6, 1, 7, 8, 9]. Recent studies have suggested that the immunopathology is virus-independent (although virus-triggered) [4, 10], and have identified central roles for the proinflammatory cytokines IL-6 and granulocyte-macrophage colony stimulating factor (GM-CSF) [5]. Furthermore, the beneficial effect of anti-inflammatory treatments (such as corticosteroids [11] and IL-6 receptor antagonists [12]) highlight the importance of inflammation in COVID-19 pathogenesis. Moreover, SARS-CoV-2 infection has been observed to be a poor inducer of interferon responses, due to the virus’ ability to evade cytosolic detectors, inhibit induction and interfere with signalling (see e.g. [13, 14] and the references therein). However, whilst type I and III interferons are potent antiviral cytokines, whose defensive potential is clear, excessive interferon activity can be harmful (for example, interferonopathies [15]), and interferon deficiencies (such as autoantibodies [16] and genetic susceptibilities [17, 18]) have been linked to severe disease. In spite of this, the precise mechanisms behind the development of severe COVID-19 are still unclear. Whilst immunological studies (focusing on peripheral blood [19], BAL fluid [6] or post-mortem investigations [4], for example) have revealed fundamental insights into SARS-CoV-2 infection and COVID-19, they are not without their limitations (e.g. lung immunology can be markedly different to that observed in peripheral blood).

Mathematical and computational models (which carry their own limitations) offer an additional perspective which can supplement experimental studies to enhance understanding. The majority of mathematical models applied to the within-host spread of an infection are ordinary differential equations (ODEs), and are variants on the well-known S-I-R model, predominately used to study transmission dynamics. The popularity of these models is a testament to their ease of construction and simulation, as well as their ability to be fitted to experimental data, and have proven to be highly informative for viral induced immunopathology (see e.g. [20]). Many of these types of models have been applied to SARS-CoV-2 infection and COVID-19 [21, 22, 23, 24], as well as other viral infections and bacterial co-infections [25, 26, 27, 28], and *in-silico* clinical trials [29]. However, ODE-type models neglect variations in space: the constraints of anatomical and physiological parameters. Individual-based models (IBMs), also known as agent-based models (ABMs), are a common computational technique to study spatially-dependent systems and have been widely used to study host-pathogen systems [30, 31, 32, 33, 34]. There are several groups applying multiscale IBMs (or ABMs) to study the within-host spread of SARS-CoV-2 infection [35, 36, 37, 38, 39].

In this article, we develop a hybrid, multiscale, individual-based model to study the within-host spread of SARS-CoV-2 and the subsequent innate immune response. We have focussed specifically on the interactions of epithelial cells, macrophages and a subset of cytokines. In Section 2, we discuss the model design (§2.1), the extracellular virus (§2.2), the cytokines and chemokines (§2.3), the viral entry and replication model (§2.4), epithelial cells (§2.5), macrophages (§2.6) and parameter estimation (§2.7). Results are presented in Section 3, where we study the influence of increasing initial viral deposition (§3.1), increasing delay in interferon secretion from epithelial cells and interferon secretion rate (§3.2), as well as variations in some macrophage parameters (§3.3), on the spread of infection. Discussion is provided in Section 4 and future work is discussed in Section 5.

## 2 Mathematical and Computational Model

The presented model extends, and adapts, an established model for the within-host spread of *Mycobacterium* tuberculosis (M. Tb) infection [33] to simulate the spread of SARS-CoV-2 infection. Section 2.1 introduces the model design, whilst Sections 2.2-2.6 introduce the model components. The model supports a uniform monolayer of lung epithelial cells (see §2.5); extracellular virus diffusion (see §2.2), as well as virus-specific entry and replication pathways (see §2.4); subsequent *innate* immune response, such as macrophages, (see §2.6); and cytokine signalling from epithelial and immune cells (see §2.3). A schematic is given in Fig. 1(a).

**Figure 1:**
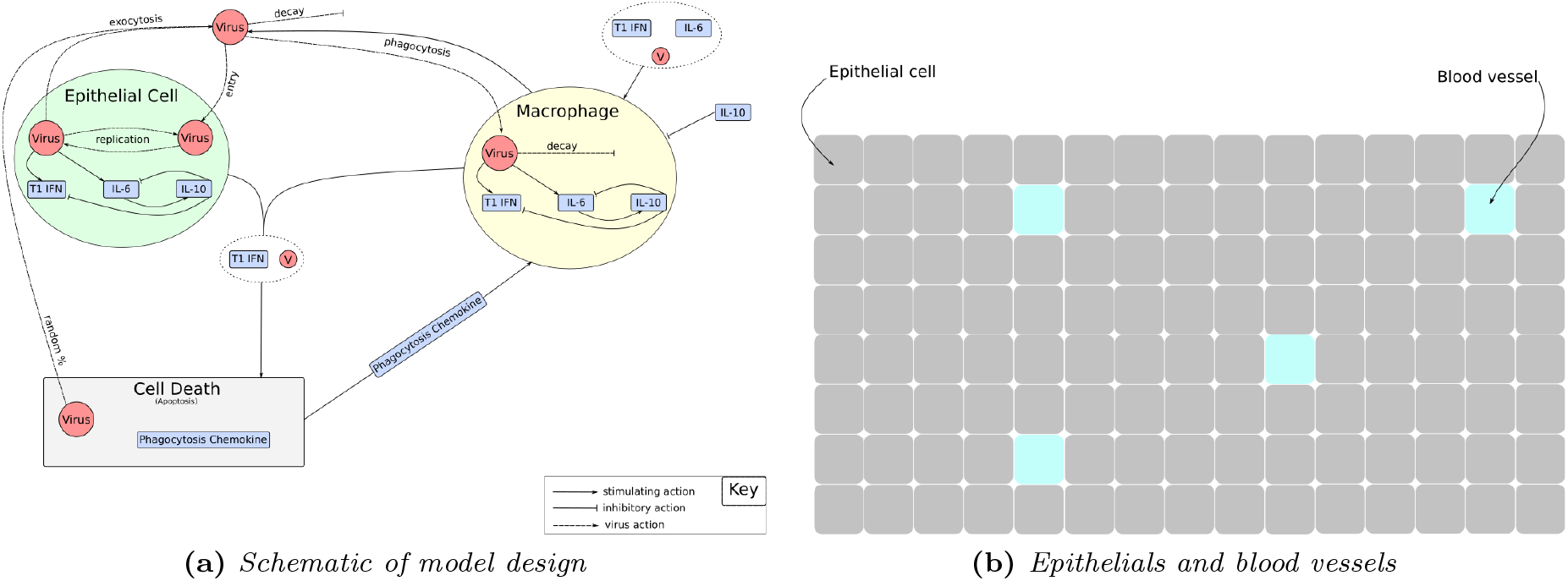
Model design.

### 2.1 Model design

Let the time interval be denoted by 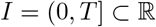 such that the closure is denoted by *Ī* = [0, *T*], where 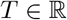 denotes the simulation time. Let 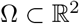 denote a two-dimensional domain containing a uniform grid of lung epithelial cells and randomly distributed cross-sections of blood vessels (see Fig. 1(b)), which is a reasonable assumption provided that the blood vessels are perpendicular to, and that there are no branching points through, the plane of interest (see Bowness *et al.* [33] and the references therein). We ignore any temporal dynamics or spatial changes of these vessels. Let Ω^*e*^ ⊆ Ω and Ω^*b*^ ⊂ Ω denote the sets of lung epithelial cells and blood vessel cross-sections, respectively, such that

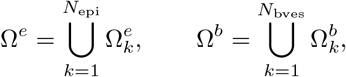

where 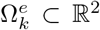, *k* = 1,…,*N*_epi_, represents a single lung epithelial cell; 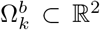, *k* = 1,…,*N*_bves_, represents the cross-section of a blood vessel; and *N*_epi_, *N*_bves_ denote the number of epithelial cells and blood vessel cross-sections, respectively. Clearly, we have Ω = Ω^*e*^ ∪ Ω^*b*^ and *N*_grid_ = *N*_epi_ + *N*_bves_, where *N*_grid_ denotes the size of the grid (see Fig. 1(b)). Furthermore, we define Ω_*T*_ = Ω × *I*, and *∂*Ω_*T*_ = Ω × *I*, where *∂*Ω denotes the boundary of Ω.

Initially, the monolayer of epithelial cells is randomly seeded with a user-defined number of individual virus particles, known as virions (see §2.2), which can be provided via a multiplicity of infection (MOI). Furthermore, the grid is initially populated with a user-defined number of *tissue-resident* immune cells (specifically, macrophages), which are assumed to be in a resting state (see §2.6).

### 2.2 Extracellular virus

Due to the high viral loads which can be observed [24, 21, 40], modelling extracellular virus particles (virions) as individual discrete agents is computationally challenging. Therefore, we choose to model the extracellular virus as a continuous field using a reaction-diffusion equation. Let 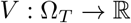 denote the *number* concentration of extracellular virus (units: virions/*mm*^2^). Then the spatiotemporal evolution of the extracellular virus is given by

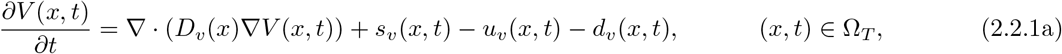

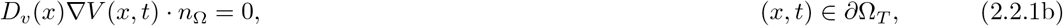

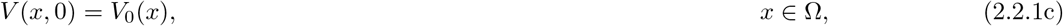

where *D_ν_* denotes the spatially-dependent viral diffusion coefficient; *s_ν_* denotes the source of virus from exocytosed virions following replication; *u_ν_* denotes the uptake of virus following engagement with ACE2 receptors; *d_ν_* denotes the decay of extracellular virus; denotes the outward unit normal to Ω from *∂*Ω; and 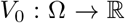 denotes the spatially-dependent initial condition, which is determined from either a user-defined number of virions or a user-defined initial multiplicity of infection (MOI). A similar PDE is used for the cytokine modelling (see §2.3). The total number of extracellular virions at time *t* ∈ *I* is given by

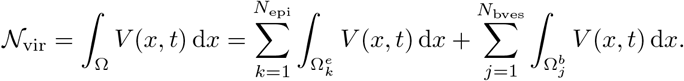

Therefore, the number of extracellular virions resident on an epithelial cell 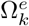, *k* = 1,…, *N*_epi_, is given by

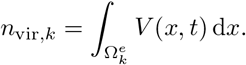

The numerical approximation to (2.2.1) is obtained using a first-order forward Euler scheme in time and a second-order central finite difference scheme in space, where each finite difference node corresponds to either a blood vessel or an epithelial cell.

### 2.3 Cytokines & Chemokines

Following infection, host-cells utilise pattern recognition receptors (PRRs) to detect pathogen associated molecular patterns (PAMPs) and damage associated molecular patterns (DAMPs), leading to the secretion of a variety of cytokines and chemokines which regulate the host response to the infection. The balance of these cytokines and chemokines is crucial in ensuring an effective and efficient management, as well as clearance, of the infection with minimal damage to the host. Indeed, elevated levels of many pro- and anti-inflammatory markers have been observed to correlate with severe COVID-19 [5, 41, 2, 1]. In particular, recent evidence suggests IL-6 and GM-CSF play a central role in severe COVID-19 [5]. Due to limits on computational resources, it is impractical for our model to consider all of the implicated cytokines and chemokines. Therefore, we choose to consider the following subset: type I interferon (IFN-I), because it induces the expression of IFN-stimulated genes (ISGs) that can inhibit viral entry and replication processes [42], has been linked to genetic susceptibility [17], and a delay in production has been suggested to increase disease severity [43, 13] as has been observed for other coronaviruses [44, 45]; interleukins (IL-6, IL-10), because of their importance in regulating inflammation [46, 47]; and a generic mononuclear phagocyte chemokine, so that macrophages may be directed towards an apoptotic cell, which needs to be removed via phagocytosis.

Similar to the model of the extracellular virus (see §2.2), we model the cytokines and chemokines as continuous fields using reaction-diffusion equations. Let 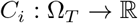, *i* = 1,…, *N*_cyt_, denote the *molar* concentration of the *i*-th cytokine (units: nanomolar (nM)), where *N*_cyt_ denotes the number of cytokines and chemokines considered in the model. Then the spatiotemporal evolution of the *i*-th cytokine or chemokine is given by

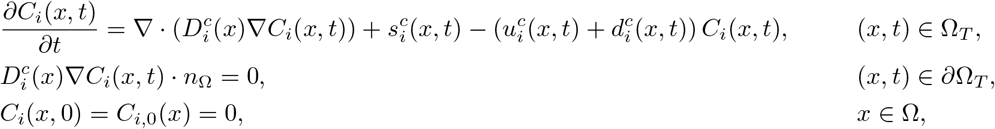

where 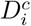 denotes the spatially-dependent diffusion coefficient; 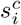 denotes the source of cytokine from host cells; 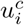 denotes the uptake of cytokine by host cells; 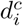 denotes the extracellular decay; *n*_Ω_ denotes the outward unit normal to Ω from *∂*Ω; and 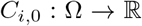 denotes the initial condition, which is set to zero. Cytokine sources from all cells satisfy:

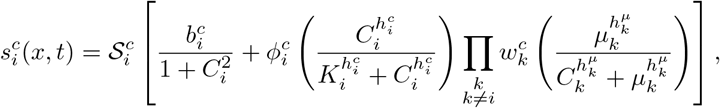

where 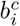 denotes the basal source; 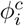 denotes the maximal production rate; *K_i_* denotes the maximal production half-max; 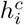 denotes the maximal production Hill coefficient; *μ_k_* denotes the inhibitory half-max; 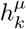 denotes the inhibitory Hill coefficient; 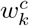 denotes the inhibitory weights; and 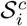 denotes the signal. Therefore, cytokines can only be produced in the presence of a sufficiently strong signal; the basal source is required to initiate the secretion, but reduces at a sufficient high level; a maximal rate is reached at sufficiently high levels; and cytokine production may be inhibited by other cytokines, but not removed entirely because of the basal production. The signal is given by

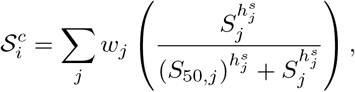

where *w_j_* denotes the signal weights; *S*_50,*j*_ denotes the signal half max; 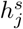 denotes the signal Hill coefficient; and *S_j_* denotes the component (such as a specific cytokine or extracellular/intracellular virus) that produces the signal. Note that each host cell will define individual sources, uptakes and signals depending on cell class and state. Reaction-diffusion parameters are given in Table 1, whilst cell-specific source, uptake and signal parameters can be found in Tables 3 and 4, respectively.

**Table 1:** Reaction-diffusion baseline parameters.

| Description | Value | Reference / Justification |
| --- | --- | --- |
| Grid spacing ($\Delta x$) | 0.02 mm | [33] |
| Cytokine diffusion coefficient ($D_i^c$) | 0.036 mm<sup>2</sup> hr<sup>-1</sup> | [33] |
| Virus diffusion coefficient ($D_v$) | 0.0054 mm<sup>2</sup> hr<sup>-1</sup> | Calculated from [36] |
| Cytokine decay coefficient ($d_i^c$) | 0.612 hr<sup>-1</sup> | [33] |
| Virus decay coefficient ($d_v$) | 0.02 hr<sup>-1</sup> | Chosen to be approximately an order of magnitude lower than the cytokine decay coefficient |

**Table 2:** Viral entry and replication baseline parameters.

| Description | Value | Reference / Justification |
| --- | --- | --- |
| Total number of receptors ($[R_{tot}]$) | $10^3 - 10^4$ | [36] |
| Binding rate ($r_{bind}$) | $0.06 \text{ hr}^{-1}$ | Calculated from [36] |
| Recycle rate ($r_{recycle}$) | $0.6 \text{ hr}^{-1}$ | Calculated from [36] |
| Endocytosis rate ($r_{endo}$) | $0.6 \text{ hr}^{-1}$ | Calculated from [36] |
| Release rate ($r_{release}$) | $0.06 \text{ hr}^{-1}$ | Calculated from [36] |
| Uncoating rate ($r_U$) | $0.6 \text{ hr}^{-1}$ | Calculated from [36] |
| RNA synthesis rate ($r_P$) | $0.6 \text{ hr}^{-1}$ | Calculated from [36, 35] |
| RNA decay rate ($\lambda_R$) | $0.021 \text{ hr}^{-1}$ | Calculated from [36] |
| Protein synthesis rate ($r_S$) | $0.6 \text{ hr}^{-1}$ | Calculated from [36] |
| Assembly rate ($r_A$) | $0.6 \text{ hr}^{-1}$ | Calculated from [36, 35] |
| Protein decay rate ($\lambda_P$) | $0.021 \text{ hr}^{-1}$ | Calculated from [36] |
| Export rate ($r_E$) | $0.6 \text{ hr}^{-1}$ | Calculated from [36, 35] |
| Maximum RNA synthesis ($r_{max}$) | $0.3 \text{ hr}^{-1}$ | Calculated from [35] |
| RNA synthesis half-max ($r_{1/2}$) | 2000 | [35] |
| Half-max value of inhibitory IFN-I ($C_{50,IFN-I}$) | $0.004 \text{ nM hr}^{-1}$ | Heuristically chosen |
| Hill coefficient of inhibitory IFN-I ($h_{IFN-I}$) | 2 | Heuristically chosen |

**Table 3:** Epithelial baseline parameters. Values which are varied in Section 3 are highlighted in a light gray colour.

| Description | Value | Reference / Justification |
| --- | --- | --- |
| Width and height of epithelial cell ($\Delta x$) | 0.02 mm | [33] |
| Healthy basal cytokine secretion ($b_i^c$) | 0.0 nM hr<sup>-1</sup> | Assumed |
| Healthy cytokine secretion ($\phi_i^c$) | 0.0 nM hr<sup>-1</sup> | Assumed |
| Healthy cytokine uptake ($u_i^c$) | 0.0 nM hr<sup>-1</sup> | Assumed |
| Infected basal cytokine secretion ($b_i^c$) | 0.0126 nM hr<sup>-1</sup> | Assumed to be the same as $\phi_i^c$ |
| Infected cytokine secretion ($\phi_i^c$) | 0.0126 nM hr<sup>-1</sup> | Calculated from [35] |
| Infected cytokine uptake ($u_i^c$) | 0 nM hr<sup>-1</sup> | Assumed |
| Cytokine half-max ($K_i$) | 0.01 nM | Heuristically estimated from [35] |
| Cytokine hill coefficient ($h_i$) | 2 | Assumed |
| Virus-dependent cytokine signal half-max ($S_{50,V}$) | 1 virion | Assumed |
| Cytokine signal hill coefficient ($h_j$) | 2 | Assumed |
| Cytokine inhibition half-max ($\mu_k$) | 5.0 nM | Chosen so that inflammatory state can be established |
| Cytokine inhibition hill coefficient ($h_k$) | 8 | Chosen so that change to anti-inflammatory state is rapid |
| Cytokine-dependent cytokine signal half-max ($S_{50,C}$) | 0.1 nM | Heuristically estimated from [35] |
| Burst threshold by virions | 10<sup>6</sup> | Chosen from volume considerations |
| Apoptosis timescale ($T_{\text{apop}}$) | 0.1 hr | Heuristically estimated from [36, 35] |
| Apoptosis by virus max rate ($r_{\text{max},[A]_k}^{\text{apop}}$) | 10<sup>-5</sup> hr<sup>-1</sup> | Heuristically chosen |
| Apoptosis by virus half-max ($[A]_{k,50}^{\text{apop}}$) | 5000 virions | Heuristically chosen |
| Apoptosis by virus hill coefficient ($n_{[A]_k}^{\text{apop}}$) | 1 | Assumed |
| Apoptosis by T1 IFN half-max ($C_{k,50,\text{IFN-I}}^{\text{apop}}$) | 1 nM | Heuristically chosen |
| Apoptosis by T1 IFN hill coefficient ($n_{\text{IFN-I}}^{\text{apop}}$) | 1 | Assumed |
| Apoptosis by T1 IFN max rate ($r_{\text{max},\text{IFN-I}}^{\text{apop}}$) | 10<sup>-5</sup> hr<sup>-1</sup> | Chosen to be the same as $r_{\text{max},[A]_k}^{\text{apop}}$ |
| Phagocytosis timescale ($T_{\text{phag}}$) | 0.1 hr | Heuristically estimated from [36] |

**Table 4:** Macrophage baseline parameters. Values which are varied in Section 3 are highlighted in a light gray colour.

| Description | Value | Reference / Justification |
| --- | --- | --- |
| Radius | 0.01 mm | [33] |
| Resting basal cytokine secretion ($b_i^c$) | 0 nM hr<sup>-1</sup> | Assumed |
| Resting cytokine secretion ($\phi_i^c$) | 0 nM hr<sup>-1</sup> | Assumed |
| Resting cytokine uptake ($u_i^c$) | 0 nM hr<sup>-1</sup> | Assumed |
| Active basal cytokine secretion ($b_i^c$) | 0.00126 nM hr<sup>-1</sup> | Assumed to be the same as $\phi_i^c$ |
| Active cytokine secretion ($\phi_i^c$) | 0.00126 nM hr<sup>-1</sup> | Calculated from [35] |
| Active cytokine uptake ($u_i^c$) | 0 nM hr<sup>-1</sup> | Assumed |
| Cytokine half-max ($K_i$) | 0.001 nM | Chosen to be an order of magnitude lower than for epithelial cells |
| Cytokine hill coefficient ($h_i$) | 2 | Assumed |
| Virus-dependent cytokine signal half-max ($S_{50,V}$) | 1 virion | Assumed |
| Cytokine signal hill coefficient ($h_j$) | 2 | Assumed |
| Cytokine inhibition half-max ($\mu_k$) | 5 nM | Chosen to be the same as epithelial cells |
| Cytokine inhibition hill coefficient ($h_k$) | 8 | Assumed |
| Cytokine-dependent cytokine signal half-max ($S_{50,C}$) | 0.01 nM | Chosen to be an order of magnitude lower than for epithelial cells |
| Resting phagocytosis rate of virus ($r_{[V]}$) | 0.006 hr<sup>-1</sup> | Chosen to be an order of magnitude lower than when active |
| Active phagocytosis rate of virus ($r_{[V]}$) | 0.06 hr<sup>-1</sup> | Chosen to be the same as $r_{\text{bind}}$ |
| Resting internalised virus decay rate ($d_{[V]}$) | 0 hr<sup>-1</sup> | Assumed |
| Active internalised virus decay rate ($d_{[V]}$) | 0.2 hr<sup>-1</sup> | Chosen to be an order of magnitude higher than the extracellular decay rate |
| Internalised virus inhibition half-max ($[V]_{50}$) | $5 \times 10^5$ virions | Chosen to be half the burst threshold |
| Internalised virus inhibition Hill coefficient ($n_{[V]}$) | 8 | Assumed |
| Resting migration rate | 0.333 hr | [33] |
| Active migration rate | 7.8 hr | [33] |
| Migration bias ($m_b$) | 3.0 | [33] |
| Phagocytosis chemokine migration weight ($w_{pck}^{\text{migrate}}$) | 1 | Assumed |
| IL-6 and phagocytosis chemokine recruitment weights ($w_C^{\text{recruit}}$) | 0.5 | Chosen to ensure equal weighting |
| IL-10 recruitment weight ($w_{\text{IL-10}}^{\text{recruit}}$) | 1 | Assumed |
| Recruitment half-max ($C_{k,50}^{\text{recruit}}$) | 0.03 nM | Heuristically chosen |
| Recruitment Hill coefficient ($n^{\text{recruit}}$) | 3 | Heuristically chosen |
| IFN-I, IL-6 and phagocytosis chemokine activation weights ($w_C^{\text{activate}}$) | 0.25 | Assumed |
| IL-10 activation weight ($w_{\text{IL-10}}^{\text{activate}}$) | 1 | Assumed |
| Activation probability ($p_{\text{activate}}$) | 0.1 | Assumed |
| Activation half-max ($C_{k,50}^{\text{activate}}$) | 0.006 nM | Heuristically estimated from [35] |
| Activation Hill coefficient ($n^{\text{activate}}$) | 8 | Assumed |
| Bursting threshold | $10^6$ | Chosen to be the same as epithelial cells |
| Apoptosis timescale ($T_{\text{apop}}$) | 0.1 hr | Chosen to be the same as epithelial cells |
| Apoptosis by IFN-I half-max ($C_{k,50,\text{IFN-I}}^{\text{apop}}$) | 0.1 nM | Chosen to be an order of magnitude lower than for epithelial cells |
| Apoptosis by IFN-I Hill coefficient $n_{\text{IFN-I}}^{\text{apop}}$ | 1 | Assumed |
| Apoptosis by IFN-I max rate $r_{\text{max,IFN-I}}^{\text{apop}}$ | $10^{-4}$ hr<sup>-1</sup> | Chosen to be an order of magnitude higher than for epithelial cells |
| Phagocytosis timescale ($T_{\text{phag}}$) | 0.1 hr | Chosen to be the same as epithelial cells |
| Phagocytose apoptotic epithelial probability | 1.0 | Assumed |
| Phagocytose infected epithelial probability | $10^{-4}$ | Chosen to ensure low probability |
| Phagocytose apoptotic immune probability | 1.0 | Assumed |

### 2.4 Viral entry and replication

SARS-CoV-2 virions consist of four main structural proteins: the spike (S) protein, which is exposed at the surface and facilitates entry into target cells; the membrane (M) and envelope (E) proteins, which are important for morphogenesis and budding; and the nucleocapsid (N) protein, which encapsidates the viral genome [48]. The S protein facilitates entry into target-host cells by binding with host-cell surface receptors; specifically, the angiotensin-converting enzyme 2 (ACE2) receptor [48]. ACE2 is expressed in a variety of human tissues, but is abundantly expressed in upper and lower respiratory tracts [49]. Following ACE2 receptor binding, the S protein is cleaved (either at the cell surface or endosome) by cellular proteases (such as TMPRSS2 [50, 51] or cathepsins [52]), enabling fusion to take place which leads to the uncoating of the viral envelope and the subsequent deposition of the viral genome to the host-cell cytoplasm.

The SARS-CoV-2 genome is a single-stranded, positive-sense RNA genome that is capped and polyadenylated, and possesses cis-acting elements necessary for controlling viral RNA replication. The replication of the full length, positive-sense, viral genomic RNA produces a full length intermediate negative-sense RNA genome, which serves as a template for the production of new full length, positive-sense, genomic RNA [48]. The newly synthesised positive-sense RNA genomes are either packaged into new virions or are used for further replication. A feature of coronavirus replication is the discontinuous transcription process that produces sub-genomic RNA (sgRNA) [53]. The sgRNA is important for the production of the structural and accessory proteins that are required for the assembly process [48]. In general, newly assembled virions are removed from the cell by exocytosis [54].

In the model presented, the viral entry and replication procedures are defined to be *surface* and *bulk* components of a *pathway,* respectively. Each epithelial cell contains its own *pathway* model, allowing individual variation of parameters between epithelial cells. The viral entry and replication procedures discussed above are complex and therefore, as a starting point, we modify a pre-existing simplified model, presented in [36], to include cytokine inhibition of both viral entry and replication. More complex models are available in the literature (see e.g. [55, 56]) and their inclusion into the presented model is a subject of future work.

For each epithelial cell 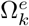, *k* = 1,…, *N*_epi_, the viral entry model, presented in [36], considers external unbound ACE2 receptors ([*R_eu_*]_*k*_), external virus-bound receptors ([*R_eb_*]_*k*_), internalised virus-bound receptors ([*R_ib_*]_*k*_) and internalised unbound receptors ([*R_iu_*]_*k*_), and is given by

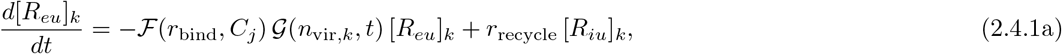

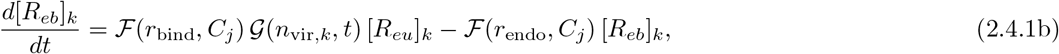

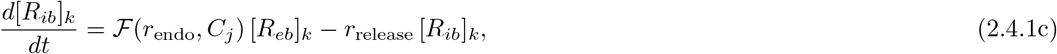

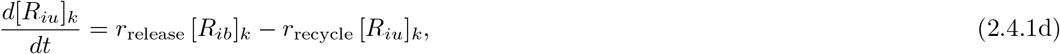

where the functions 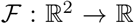 and 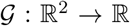 describe the rate reduction caused by an inhibiting cytokine (such as type I interferon) and whether the virus should be internalised, and are given by

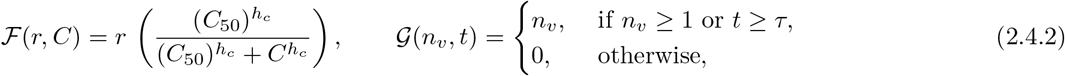

where *r* denotes the rate to be reduced; *C* denotes the inhibiting cytokine; *h_c_* denotes the Hill coefficient; *C*_50_ denotes the half-max value of the inhibiting cytokine; *n_ν_* denotes the number of virions; and *τ* denotes the time at which *n_ν_* first exceeds one. Therefore, the function 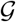 is a blocking function which prevents continuous internalisation of virions until a single whole virion is detected. Note that the blocking function is required to prevent spurious spread of infection through the epithelial monolayer at low levels of extracellular virus.

Similarly, for each epithelial cell 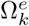, *k* = 1,…, *N*_epi_, the viral replication model, presented in [36], considers fully endocytosed virions ([*V*]_*k*_), uncoated viral RNA ([*U*]_*k*_), synthesised viral RNA ([*R*]_*k*_), synthesised viral proteins ([*P*]_*k*_) and fully assembled virions ([*A*]_*k*_), and is given by

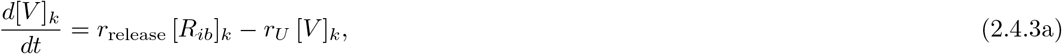

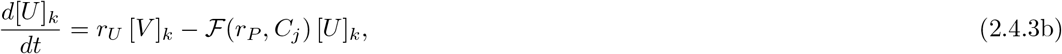

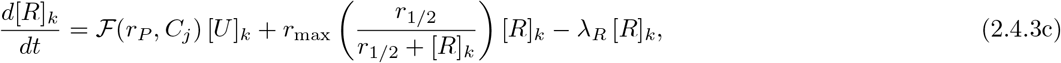

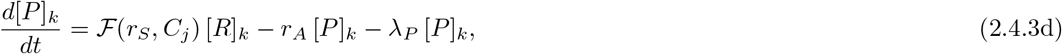

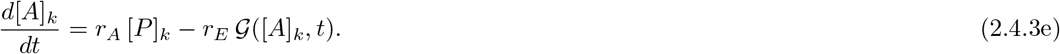

where exocytosis of newly assembled virions occurs once a single newly assembled virion is detected. Therefore, the source and uptake of extracellular virus is given by

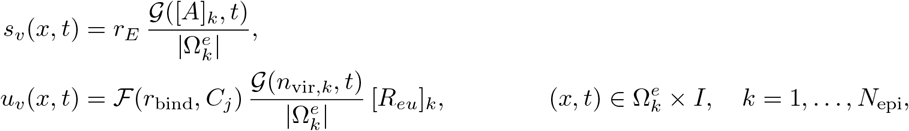

where 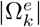 denotes the area (or volume) of 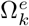.

Figure 2 illustrates the viral entry (2.4.1) and replication (2.4.3) dynamics of a single epithelial cell infected by a single virion in the absence of viral diffusion (*D_ν_* = 0), and compares against the solution of the ODE systems^1^ given by (2.4.1) and (2.4.3). Clearly, excellent agreement is seen over both short and long timescales for both entry and replication dynamics. Viral entry and replication parameters are given in Table 2. The numerical solution of (2.4.1) and (2.4.3) is obtained using a fourth-order Runge-Kutta scheme, which is available as part of the GNU Scientific Library [57]. Note that we have not observed any numerical instabilities due to discontinuous export.

**Figure 2:**
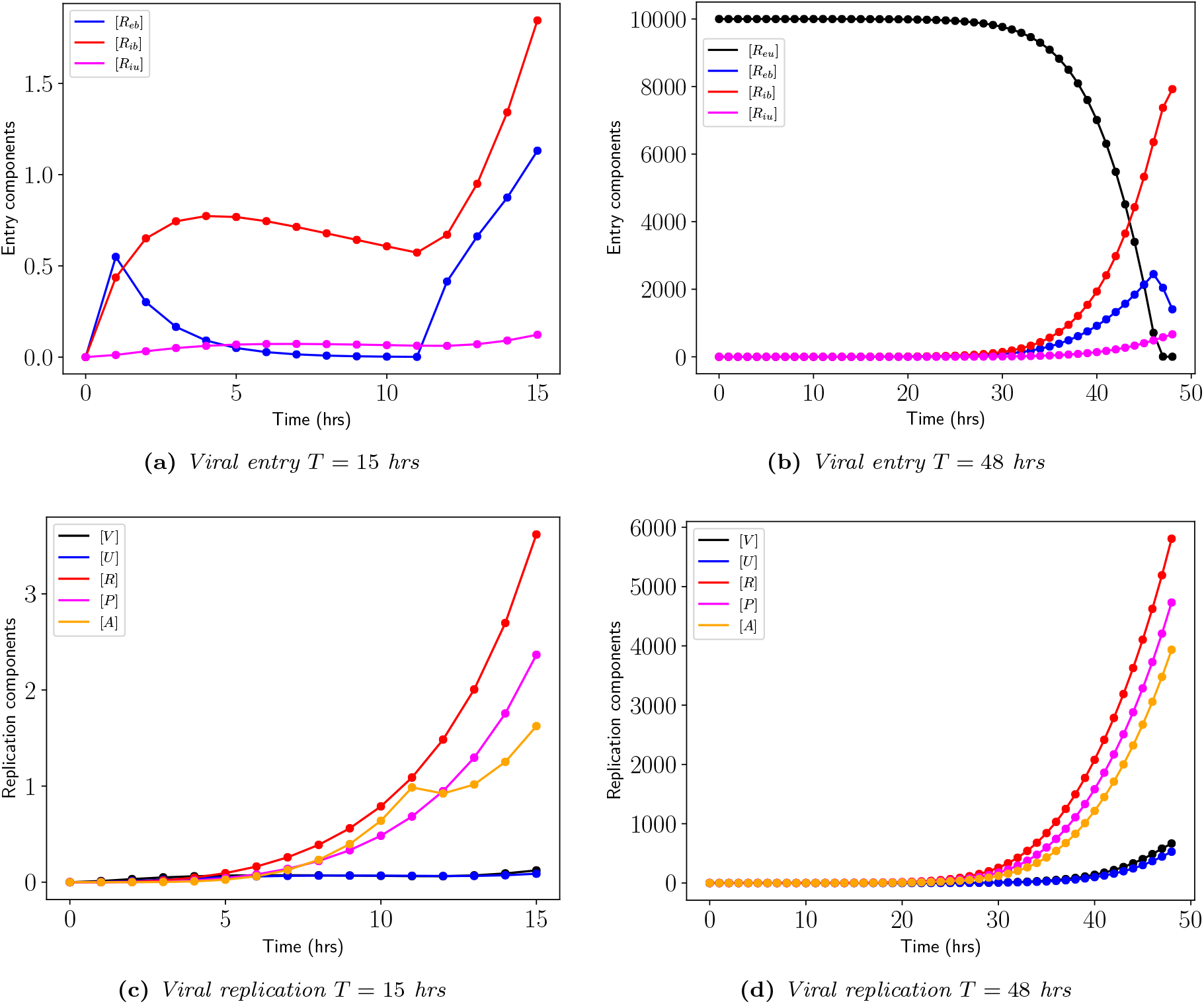
(Validation of viral entry and replication) *Comparison between presented model in the absence of diffusion (dots) and the ODE systems (solid lines), given by* (2.4.1) *and* (2.4.3), *for simulation times (a), (c) T* =15 *and (b), (d) T* = 48 *hours, respectively. Baseline parameter values are given in Table 2*.

### 2.5 Epithelial cells

As discussed in §2.1, the grid of epithelial cells Ω^*e*^ is assumed to be uniform and therefore, each epithelial cell is chosen to be a square with width and height of 20 *μ*m. Each epithelial cell can exist in five possible states: healthy, infected, infectious, apoptotic and removed. A healthy epithelial cell is placed in an infected state as soon as the number of internal bound receptors are non-zero ([*R_ib_*]_*k*_ > 0, *k* = 1,…, *N*_epi_). Therefore, an infected epithelial cell is an epithelial cell which has been infected by the virus but is not yet virus-producing (infectious). Due to the blocking function (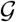 in (2.4.1)), an infected epithelial cell becomes infectious when a single whole virion is detected on the inside of the cell. As discussed in §2.4, each epithelial cell has a copy of the viral entry (2.4.1) and replication (2.4.3) models. An infectious epithelial cell may be placed into an apoptotic state due to either high levels of type 1 interferon (IFN-I) or high levels of intracellular, newly assembled, virions ([*A*]_*k*_). In both cases, a Hill function 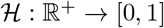 is used to determine the likelihood of triggering apoptosis in an epithelial cell, and is given by

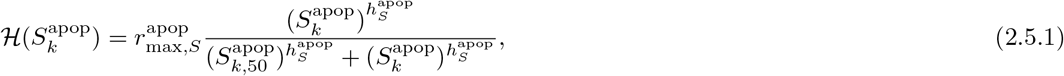

where 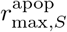 denotes the maximum apoptosis rate for signal *S*; 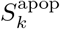 denotes the apoptosis signal (such as IFN-I or [*A*]_*k*_) for the *k*-th epithelial cell; 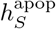 denotes the Hill coefficient for signal *S*; and 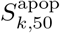 denotes the half-max value. An epithelial cell will then become apoptotic provided 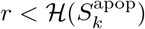, where *r* ∈ [0,1] is a uniform random number. Apoptotic epithelial cells are targeted for phagocytosis by active macrophages (see §2.6.4), and are therefore placed into a removed state upon successful phagocytosis.

We do not attempt to model homeostasis and therefore, healthy epithelial cells do not secrete cytokines. However, once a cell is infected, we assume that it produces cytokines. For computational reasons, we assume that all cytokines considered in the model can be produced by infected epithelial cells, according to the procedure detailed in §2.3, but in varying quantities (see Table 3). As discussed in §2.3, each cytokine produced requires a specific signal with specific parameters for that cytokine. For each cytokine produced by infected epithelial cells, the signal is defined as the amount of intracellular, newly assembled, virions, with the exception of IL-10 whose signal is the level of IL-6. Epithelial cells in the apoptotic state can only produce the generic mononuclear phagocyte chemokine, which chemotactically directs macrophages to the apoptotic epithelial cell so that phagocytosis may take place. Epithelial cells in a removed state cannot produce any cytokines. Additionally, the diffusion coefficients of cytokines and extracellular virus at the location of a removed epithelial cell are set to zero, to simulate an empty space on the grid.

Figure 3(a) illustrates the spread of infection in the absence of immune cells. Light blue dots depict infected epithelial cells (not virus producing), dark blue dots depict infectious epithelial cells (virus producing) and grey dots depict apoptotic epithelial cells. Initially a single virion is placed on a single epithelial cell in the centre of the domain. As expected, the single virion infects the epithelial cell upon which it is resident, replicates inside the epithelial cell to produce new virions which infect neighbouring epithelial cells, resulting in a uniform spread of infection (as in this example, each epithelial cell is assumed to have the same number of ACE2 receptors). As can be seen from Fig. 3(a), infected (but not virus producing) epithelial cells form an outer ring of the spread, whilst infectious epithelial cells are concentrated in the centre of the spread. Apoptotic epithelial cells may be seen in both the centre of the spread, and away from the site of infection. The epithelial cells away from the site of infection are placed into an apoptotic state due to IFN-I levels. Figure 3(b) shows levels of IFN-I, normalised by the maximum value. As expected, the highest levels of IFN-I are found at the site of infection. Baseline parameter values may be found in Table 3.

**Figure 3:**
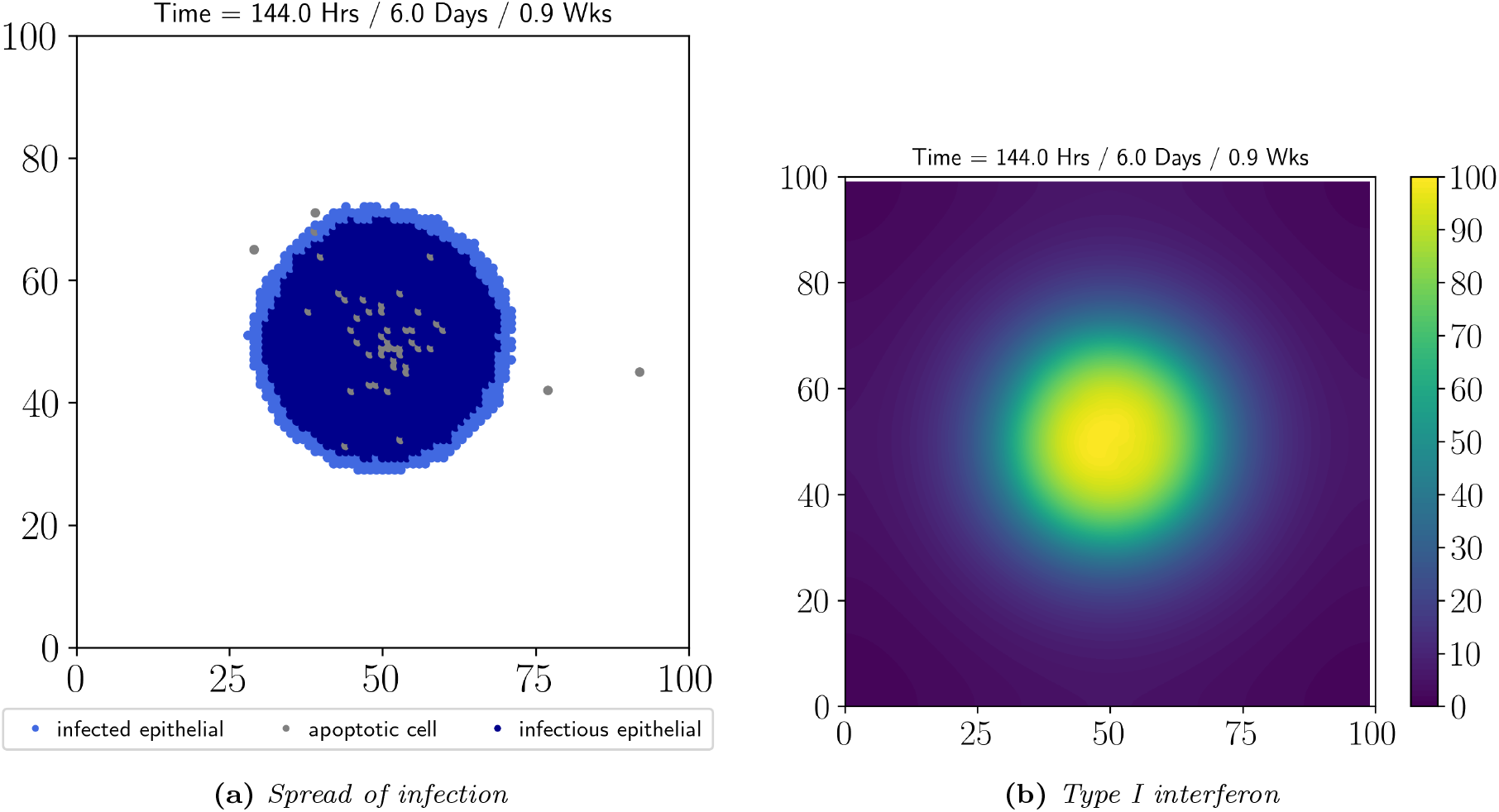
(Spread of infection) *Illustration of (a) the spread of infection through an epithelial monolayer and (b) type I interferon levels. Baseline parameters may be found in Table 3*.

### 2.6 Immune cells

Although the model presented supports macrophages, monocytes, neutrophils, and natural killer cells, we choose to focus only on macrophages for the purposes of this article. The reason for this choice, is primarily due to model complexity, but is also informed by mounting evidence suggesting an important role for macrophages [58, 59]. Evidence suggests that there is an early loss of resident alveolar macrophages, as well as a recruitment of inflammatory interstitial and monocyte-derived macrophages (see e.g. [60] and the references therein). However, for our initial model, we assume that for each class of immune cell (such as a macrophage) there is only a single type (such as alveolar macrophage) and we do not distinguish between phenotypes. We recognise the clear limitation to this assumption, and extending the model to incorporate different types and phenotypes is a subject of future work. Immune cells may, therefore, exist in three possible states: resting, active and apoptotic. The transition from resting to active state, and vice-versa, is discussed in §2.6.3. Similar to an epithelial cell, we assume an immune cell may be placed in an apoptotic state by high levels of IFN-I, or by high levels of intracellular virions. Apoptotic immune cells may be removed from the grid following successful phagocytosis by phagocytic immune cells (such as macrophages).

Again, as we do not attempt to model homeostasis, resting immune cells do not secrete cytokines. However, once an immune cell becomes active, then cytokines are secreted. Similar to epithelial cells, for computational reasons we assume that all cytokines considered in the model can be produced by immune cells, according to the procedure detailed in §2.3, but in varying quantities (see Table 4). Therefore, each immune cell is considered a major source of cytokines, but not the only source. As discussed in §2.3, each cytokine produced requires a specific signal with specific parameters for that cytokine and this is discussed in more detail in §2.6.4. Furthermore, we assume that immune cells in the apoptotic state cannot produce any cytokines, nor can they migrate.

#### 2.6.1 Migration

Each immune cell can migrate from epithelial cell to epithelial cell and is only allowed to reside on a single epithelial cell at any one time (lattice-based movement across finite difference grid). The migration may be pseudo-random or may be directed in response to cytokine levels. Each immune cell migrates according to a characteristic movement rate, which is allowed to vary for each immune cell (see Table 4). Note that we assume that apoptotic immune cells cannot migrate.

Let 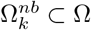 denote the set of immediate neighbours of epithelial cell 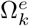, *k* = 1,…, *N*_epi_, such that

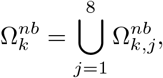

where 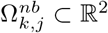, *j* = 1,…, 8, can be either an epithelial cell or a blood vessel cross-section. Note that we do not allow immune cells to migrate onto blood vessels. If an immune cell is resident on the epithelial cell 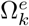, *k* = 1,…, *N*_epi_, then for each neighbour 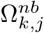, *j* = 1,…, 8, we calculate a *migrating* signal 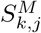, which is defined as the weighted sum of chemokines on the *j*-th neighbour of the *k*-th epithelial cell. Note that the chemokines involved are potentially specific for each immune cell, and the weights enable a migration bias towards certain chemokines for a particular immune cell. For each immune cell, we then define the total migrating signal from the immediate neighbours of the *k*-th epithelial cell as

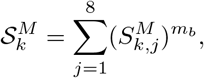

where the superscript (·)*^m_b_^* denotes a *movement bias,* which is allowed to vary for each immune cell. If the total migrating signal from the immediate neighbours is zero 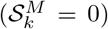, or if the immune cell state is resting, then the immune cell migrates randomly. Otherwise, the immune cell will migrate chemotactically according to the following procedure: let 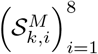 denote the sequence of partial sums given by

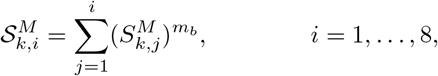

and let *r* ∈ [0,1] denote a uniform random number; then for each *i* = 1,…, 8, an immune cell which is resident on the *k*-th epithelial cell will migrate to the *i*-th neighbour provided 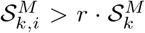 and there is sufficient space available on the *i*-th neighbour. Note that the available space is determined by the difference between the area of the *i*-th neighbour and the sum of the area of any immune cells that are resident on the *i*-th neighbour.

The pseudo-random migration of the immune cells is illustrated in Fig. 4(a), where the immune cell movement is largely clustered around its initial location before moving away. Chemotactic migration is illustrated in Fig. 4(b), where directed migration of the immune cell towards the signal (Fig. 4(d)) can be seen.

**Figure 4:**
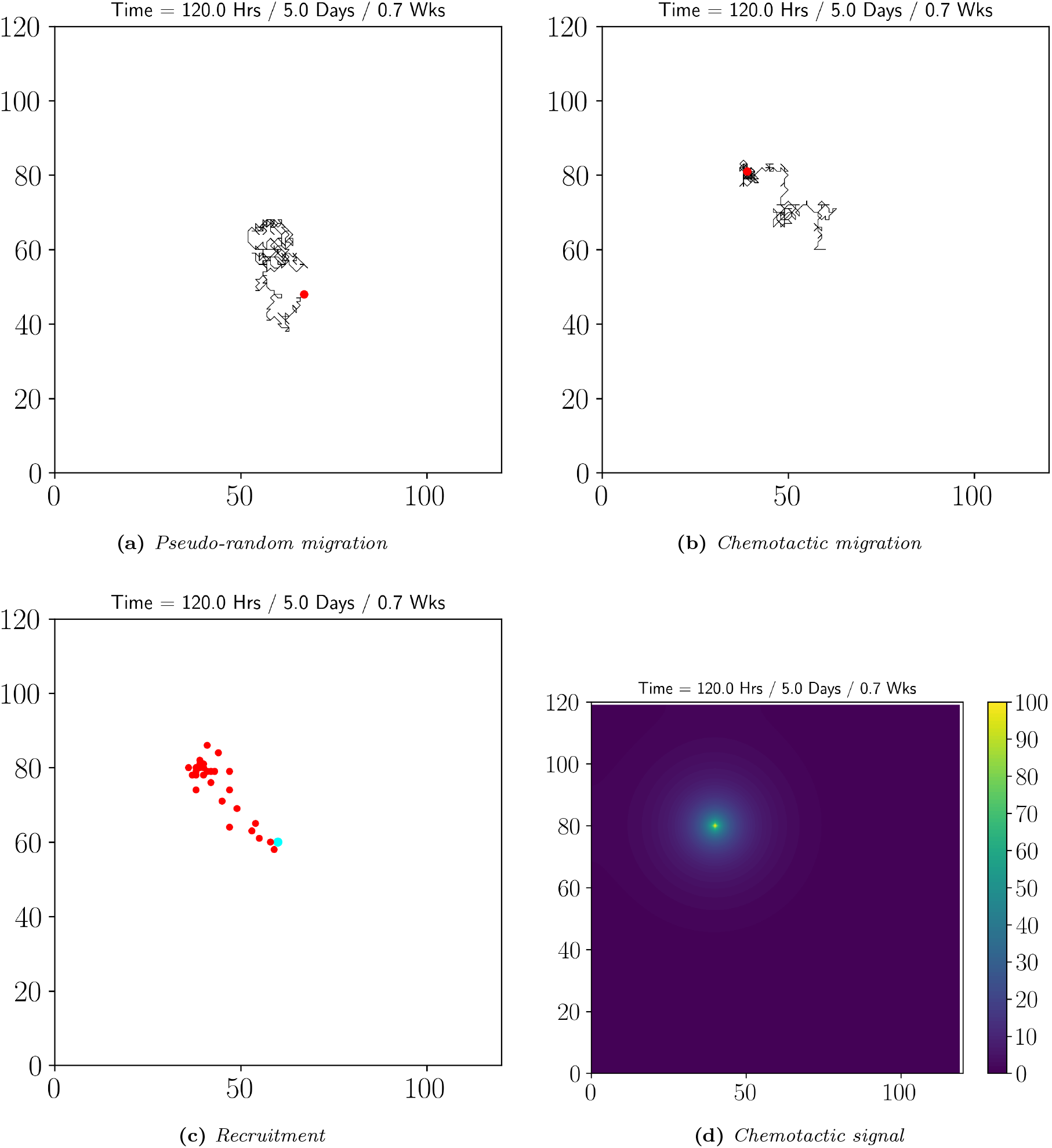
(Migration and Recruitment of Immune Cells) *Illustration of (a) pseudo-random migration of immune cells, as well as their (b) chemotactic migration and (c) recruitment, towards (d) a chemotactic signal. Note that a grid size of* 120 × 120 *has been used; a red dot denotes any immune cell; a cyan dot denotes a blood vessel; and a black line denotes an immune cell path.*

#### 2.6.2 Recruitment

The immune cells are recruited through the blood vessels, where, for each *t* ∈ *I*, each blood vessel is checked to see whether a single immune cell of each available class (for example, a macrophage) needs to be recruited from that vessel. For each neighbour of the *k*-th blood vessel, we calculate a *recruitment* signal 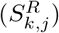, which is defined as the difference between the weighted sum of pro-inflammatory cytokines and the weighted sum of anti-inflammatory cytokines on the *j*-th neighbour of the *k*-th blood vessel. Similar to the migration procedure above, the cytokines involved are potentially specific for each immune cell, and the weights enable a recruitment bias towards certain cytokines for a particular immune cell. The total recruitment signal from the immediate neighbours of the *k*-th blood vessel is given by

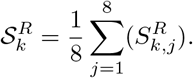

The Hill function 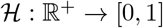 given in (2.5.1) is then used to determine the likelihood that an immune cell will be recruited from the *k*-th blood vessel. An immune cell will be recruited if 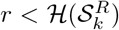, where *r* ∈ [0,1] is a uniform random number, and if there is sufficient space available on a randomly chosen neighbour of the *k*-th blood vessel. Note that if the recruitment signal is negative, then an immune cell is not recruited. All immune cells are recruited in a resting state and cannot be recruited onto a blood vessel.

#### Remark 1.

*We note that the recruitment procedure is sensitive to the choice of Hill function parameters. If the recruitment half-max, or the Hill coefficient, are too low then immune cell recruitment may be seen early in the simulation, which is undesirable as we expect the resident immune cells to be the only respondants during the early stages of infection. However, if the recruitment half-max is too high, then we may not observe any recruitment. Furthermore, if the Hill coefficient is too large, then switch-like recruitment may take place, where no recruitment is seen until the half-max, followed by saturating recruitment soon after, leading to a flooding of immune cells which can significantly affect computational time. Therefore, the Hill function parameters require manual tuning for each set of simulation parameters. Reducing this parameter sensitivity is a subject of future work.*

Figure 4(c) illustrates the recruitment of immune cells, where following a sufficiently strong chemotactic signal (Fig. 4(d)) immune cells enter the domain from a nearby vessel, before chemotactically migrating towards the signal, as expected.

#### 2.6.3 Activation & Deactivation

The activation and deactivation of the immune cells is carried out in two ways: according to local cytokine levels; and probabilistically if the extracellular/intracellular viral load is sufficient. Specifically for macrophages, if the internalised viral content exceeds one, then the macrophage will activate provided *r* × *P*_activate_, where *r* ∈ [0,1] is a uniform random number and *p*_activate_ is the probability of activation. In practice, many cytokines may be involved in the activation of the immune cells. Consequently, an immune cell which is resident on the *k*-th epithelial cell has an *activating* signal 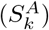, which is defined as the difference between the weighted sum of pro-inflammatory cytokines and the weighted sum of anti-inflammatory cytokines, where the cytokines involved are potentially specific for a given immune cell, similar to the migration and recruitment procedures above. Once again, the weights enable an activation and deactivation bias towards certain cytokines for a particular immune cell. The Hill function 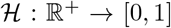 given in (2.5.1) is then used to determine the likelihood that an immune cell resident on the *k*-th epithelial cell will activate or deactivate. Therefore, an immune cell will activate if 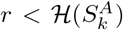, where *r* ∈ [0,1] is a uniform random number. Note that a macrophage will deactivate provided 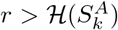, *r*_2_ > *p*_activate_ and the internalised viral content is less than one, where *r,r*_2_ ∈ [0,1] are uniform random numbers.

#### 2.6.4 Macrophages

Each macrophage is assumed to be a circle, with a diameter of 20 *μ*m. Initially, a user-defined number of resident macrophages are randomly placed in the tissue environment. Aside from the secretion of cytokines, the main role of macrophages is to patrol tissue environments and clear away any dead tissue and cells, as well as extracellular debris and pathogens [61]. In our model, both resting and active macrophages may internalise extracellular virus, whilst only active macrophages may phagocytose apoptotic epithelial and immune cells. Additionally, to aid viral clearance, we assume that an active macrophage may phagocytose an infected epithelial cell by implicitly triggering apoptosis, however this occurs infrequently in our model. As mentioned at the beginning of §2.6, we do not attempt to model homeostasis and therefore, only active macrophages produce cytokines. We assume that active macrophages produce all the cytokines considered in the model, but at varying quantities. As discussed in §2.3, cytokine production requires a specific signal with specific parameters for that cytokine. For the cytokines IFN-I and IL-6, the signal is defined as the internalised viral content, whereas for the cytokine IL-10, the signal is defined as the pro-inflammatory cytokine IL-6.

The internalisation of extracellular virus is governed by

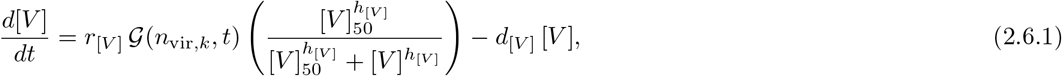

where [*V*] denotes the internalised virus; *n*_vir,*k*_ denotes the number of virions on the *k*-th epithelial cell (where the macrophage resides); 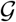 denotes the *blocking* function given in (2.4.1); *r*_[*V*]_ denotes the phagocytosis rate; [*V*]_50_ denotes the half-max value of internalised virus inhibition; *h*_[*V*]_ denotes the Hill coefficient; and *d*_[*V*]_ denotes the lysing decay of internalised virus. Parameters may be found in Table 4. Note that the parameter values are state-dependent; for example, we assume that only active macrophages breakdown internalised virus, leading to different decay values for the resting and active states. If the macrophage’s internalised viral content exceeds a user-defined threshold (see Table 4), then the macrophage may rupture, releasing the viral contents into the extracellular environment.

Phagocytosis of an infected or apoptotic epithelial cell, as well as apoptotic immune cells, is assumed to take place over a time scale which is set by the target cell (for example, an apoptotic epithelial cell). Once the phagocytosis process has begun, the target cell is placed into a temporary *phagocytosed* state, where all cellular processes stop. Additionally, the macrophage performing the phagocytosis enters a temporary *phagocytosing* state, where cytokines may still be produced but the cell cannot change state or migrate until the phagocytosis process is complete. Once phagocytosis is complete, the target cell may be placed into a removed state (in the case of epithelial cells) or it is removed from the grid (in the case of immune cells).

Resting macrophages may only migrate pseudo-randomly, whilst active macrophages may migrate chemotactically, as well as pseudo-randomly, according to the procedure detailed in §2.6.1. The chemotactic migration of macrophages is controlled solely by a generic mononuclear phagocyte chemokine, which is released by apoptotic cells (both epithelial and immune). Recruitment of resting macrophages from the blood vessels is carried out according to the procedure detailed in §2.6.2. Recruitment of the macrophages is encouraged by: IL-6; and the generic mononuclear phagocyte chemokine, whilst the anti-inflammatory cytokine IL-10 discourages recruitment.

Activation and deactivation of a macrophage is carried out according to the procedure detailed in §2.6.3. Activation of resting macrophages is encouraged by: internalised viral content [*V*]; IFN-I; IL-6; and the generic mononuclear phagocyte chemokine, whilst the anti-inflammatory cytokine IL-10 discourages the pro-inflammatory activation. Therefore, we are only considering classical activation of the macrophages in our model. All cytokines involved in the activation and deactivation of macrophages are obtained from the location where the macrophage resides. Similar to phagocytosis, apoptosis of a macrophage is assumed to take place over a time scale which is set by the macrophage. Once the apoptosis process has begun, the macrophage is placed into a temporary *apoptosed* state, where we assume that cytokines may still be released but the cell cannot change state, migrate or carry out phagocytosis; this is a reasonable approximation, particularly during the early stages of the apoptotic programme. Once apoptosis has been completed, the macrophage is placed into a apoptotic state, where the generic mononuclear phagocyte chemokine is released to encourage efferocytosis, but all other cytokine secretions stop, and the macrophage may not migrate nor carry out phagocytosis. Additionally, once apoptosis is completed, we assume that a random portion of the macrophage’s internalised viral content is scattered to the neighbouring epithelial cells.

### 2.7 Parameter estimation

Some of the parameter values, which are given in Tables 1-4, have been obtained from the modelling literature: namely, Bowness *et al.* [33], Sego *et al.* [35], and Getz *et al.* [36], which in turn obtained their parameters either heuristically, from other modelling work, or by estimation, approximation and calibration from experimental studies. Therefore, we do not claim all of the parameters are specific to SARS-CoV-2, but can be from related viruses such as SARS-CoV-1. Furthermore, not all parameter values could be directly related. Therefore, the reader will notice that specific language is used when justifying a parameter value; specifically, *assumed, chosen, heuristically chosen, heuristically estimated from* and *calculated from*. Parameter values which are *chosen* have not been heuristically varied, instead they are chosen for a particular reason or to simulate a specific behaviour; *heuristically chosen* parameters are those which have been chosen from computational experiments; *heuristically estimated from* denotes parameters which were obtained from the literature but then varied heuristically for our model; and *calculated from* denotes parameters which were obtained from the literature but altered, due to a change in units, for example.

### 2.8 Code and data availability

The code used in this article can be found on GitHub:

https://github.com/Ruth-Bowness-Group/CAModel.

The data used in this article has been archived within the Zenodo repository:

https://doi.org/10.5281/zenodo.6514656.

## 3 Results

### 3.1 Influence of initial viral deposition

To investigate the influence of the initial viral deposition, we consider three values for the initial multiplicity of infection (MOI = 0.001 (= 11 virions), 0.01 (= 102 virions), 0.1 (= 1015 virions)). Due to the possibility of the immune response being influenced by the grid size, each initial viral deposition is randomly distributed over a subdomain located in the centre of the tissue environment; the full grid size is 200 × 200, whilst the central subdomain has a grid size 100 × 100, which leaves a *buffer* around the initial subdomain for the infection to spread into. Therefore, the number of virions to be deposited is calculated using the initial MOI and the number of epithelial cells in the subdomain. Rigorous investigation into the grid size dependency is a subject of future work.

Figure 5 shows the spread of infection at the end of the simulation (T = 120 hours) as the MOI is increased (baseline parameter values are given in Tables 1-4). In Fig. 5(a), we can clearly see non-uniform spread of infection which is caused by each epithelial cell possessing a different number of ACE2 receptors. Infected (not virus producing) epithelial cells form a thin boundary around infectious (virus producing) epithelial cells. We see a high number of macrophages in either a resting, active or apoptotic state. Furthermore, we observe that the majority of macrophages at the site of infection are either in an active or apoptotic state, whilst the majority of resting macrophages are either local to the boundary of infection or away from it. As the MOI is increased (Figs. 5(b)-(c)), we see that the spread of infection increases, as well as increased number of active macrophages and, in particular, apoptotic macrophages. For MOIs = 0.01, 0.1 (Figs. 5(b),5(c)), we see an almost square spread of infection due to the high viral deposition for the number of epithelial cells in the initial subdomain.

**Figure 5:**
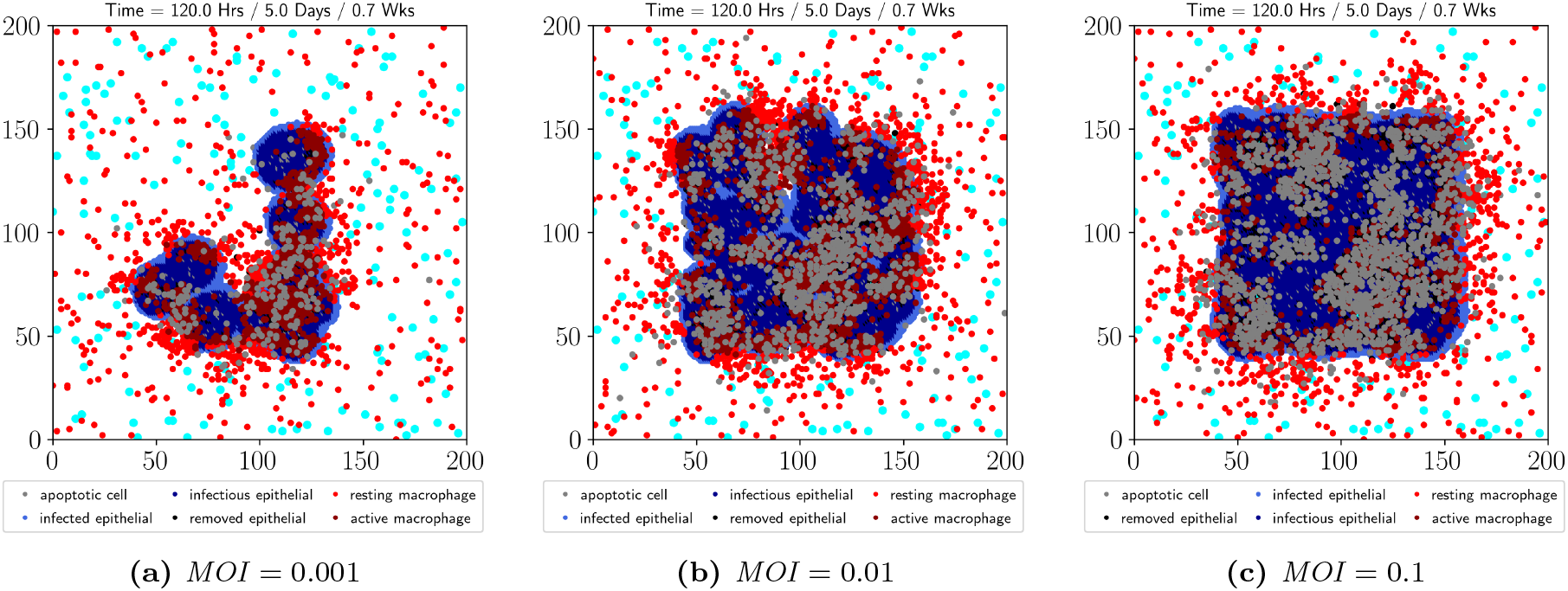
(Spread of infection for increasing MOI) *Illustration of the spread of infection at the end of the simulation (*T = 120 *hours) for (a) MOI* = 0.001, *(b) MOI* = 0.01, *and (c) MOI* = 0.1. *Baseline parameter values are given in Tables 1-4.*

Figures 6(a)-(c) illustrate the number of epithelial cells in an infectious, apoptotic or removed state, respectively, for each MOI. In Fig. 6(a), we see an increasing number of infectious epithelial cells by the end of the simulation, for increasing MOI. Additionally, there is an initial delay (≈ 12 hours) before infectious epithelial cells are first found; this initial delay is seen for all MOI values but is most easily observed for MOI = 0.1, and is a consequence of the timescale of viral entry and replication, as well as the blocking function (see §2.4), which allows a whole virion to be produced before being exocytosed. Interestingly, following the initial delay, we observe a second, longer delay before a continuous increase in infectious epithelial cells is observed, which is likely to be caused by both the blocking function and IFN-I inhibition (see §2.4 and Fig. 8(a)). Epithelial cells may be placed into an apoptotic state due to high intracellular viral loads or due to high IFN-I; in Fig. 6(b), we see an increase in apoptotic epithelial cells after *t* = 24, where, for each MOI, the start of the increase correlates with both an increase in total (across all epithelial cells) intracellular viral load (Fig. 7(a)) and extracellular IFN-I (Fig. 8(a)). Furthermore, we observe an increase in the number of apoptotic epithelial cells with increasing MOI. Epithelial cells may be placed into a removed state after successful macrophage phagocytosis; in Fig. 6(c) we see an increase in removed epithelial cells after *t* = 48 hours, which correlates with an increase in active macrophage numbers (Fig. 6(e)) for each MOI considered. Interestingly, for the largest MOI considered, we see a spike in resting macrophages at around t = 48 hours, which is not observed for the other MOIs (Fig. 6(d)), nor for active and apoptotic macrophage numbers (Figs. 6(e),(f)), and may be caused by the sharper increase in apoptotic epithelial cells at around *t* = 48 hours (Fig. 6(b)), as well as extracellular IL-6 levels (Fig. 8(b)). Alongside the increase in apoptotic epithelial cells, we see an increase in apoptotic macrophages (Fig. 6(f)), which in conjunction with increases in IL-6 levels, likely contributes to the recruitment of resting macrophages observed in Fig. 6(d). Moreover, we observe a decline in resting macrophage numbers towards the end of the simulation (Fig. 6(d)), which is likely caused by increased activation as well as increasing IL-10 levels (Fig. 8(c)), which discourages both recruitment and activation. Generally, we see an increase in resting, active and apoptotic macrophages with increasing MOI.

**Figure 6:**
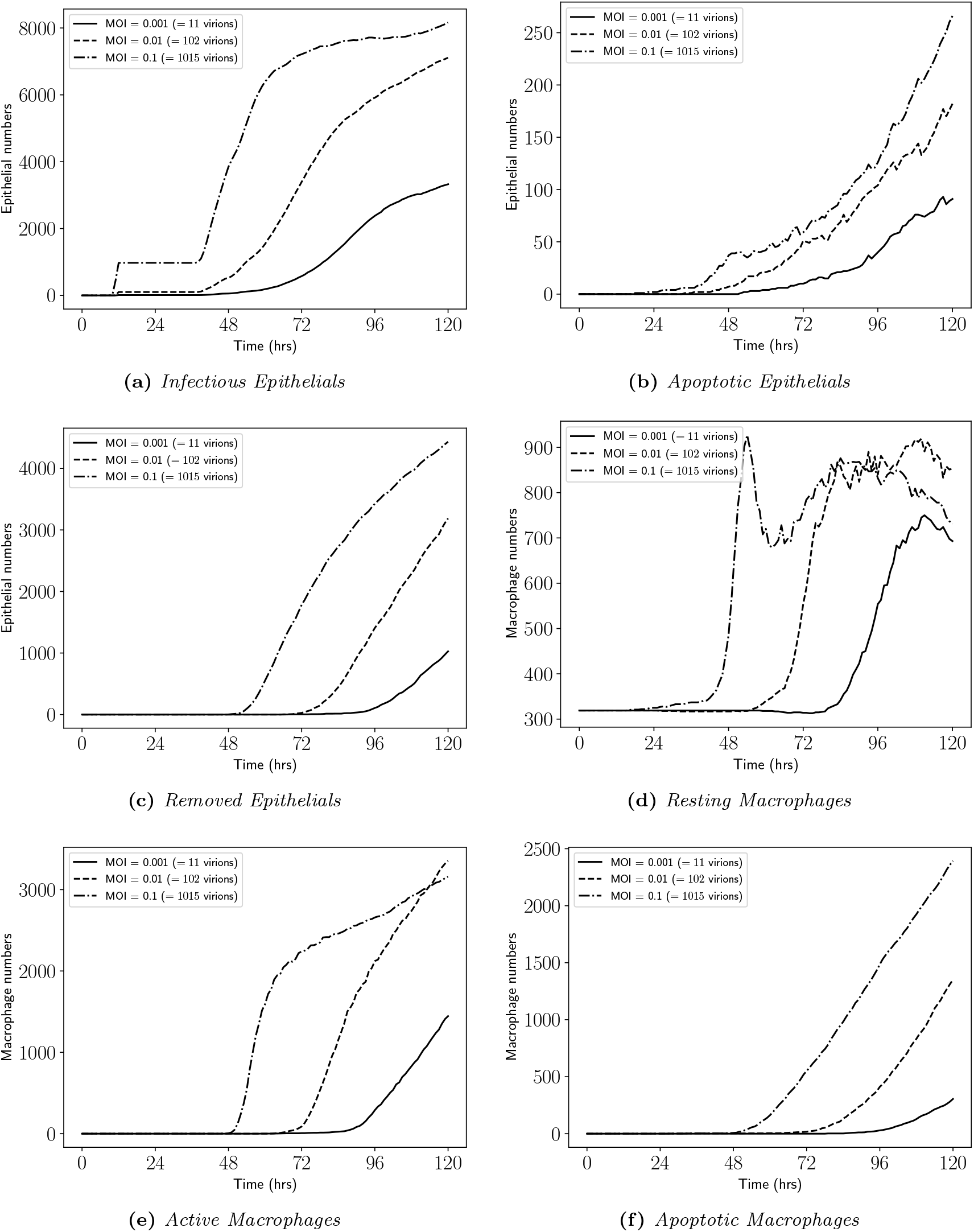
(Cell numbers for increasing MOI) *Number of (a) infectious, (b) apoptotic and (c) removed epithelial cells, as well as (d) resting, (e) active and (f) apoptotic macrophage numbers, as the simulation progresses, for increasing MOI. Baseline parameters are given in Tables 1-4*.

**Figure 7:**
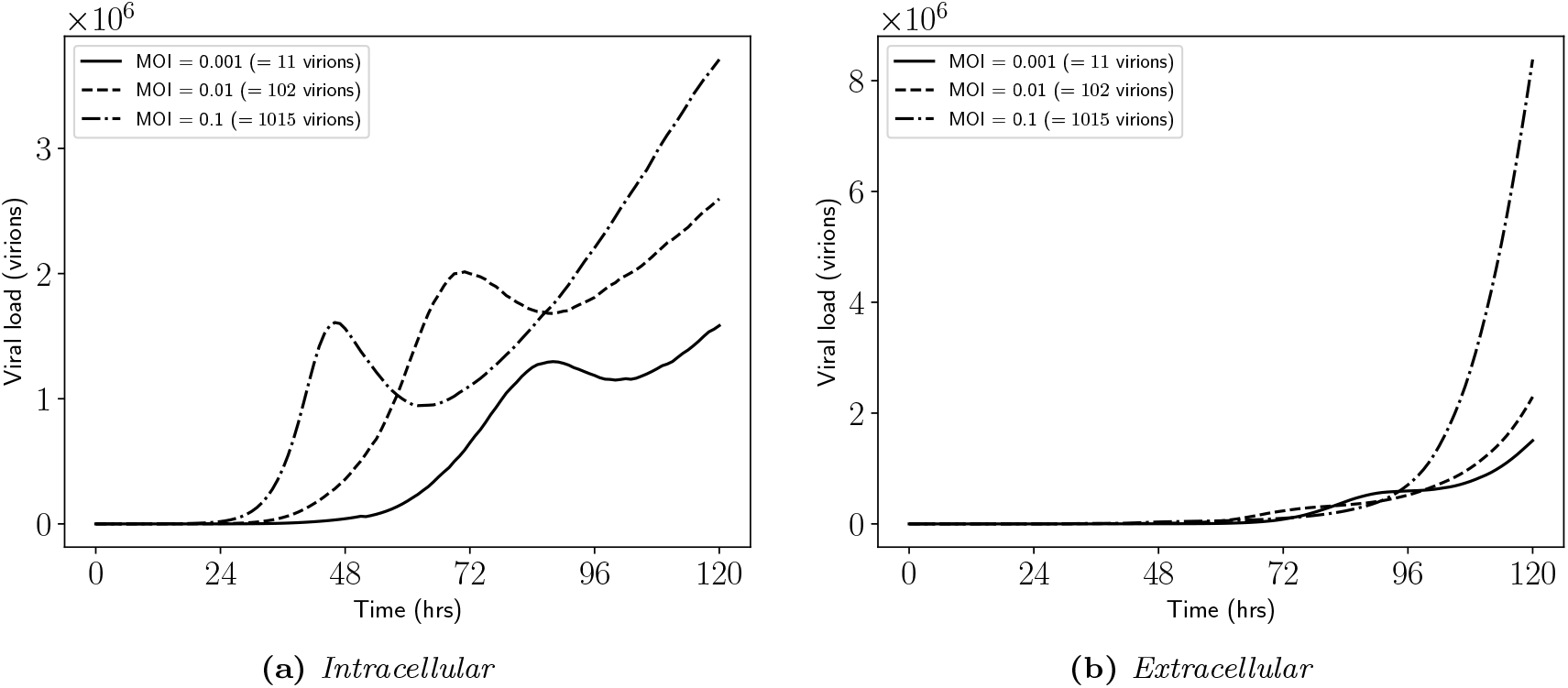
(Viral loads per grid for increasing MOI) *Illustration of (a) intracellular and (b) extracellular viral loads per grid, as the simulation progresses, for increasing MOI. Baseline parameters are given in Tables 1-4*.

**Figure 8:**
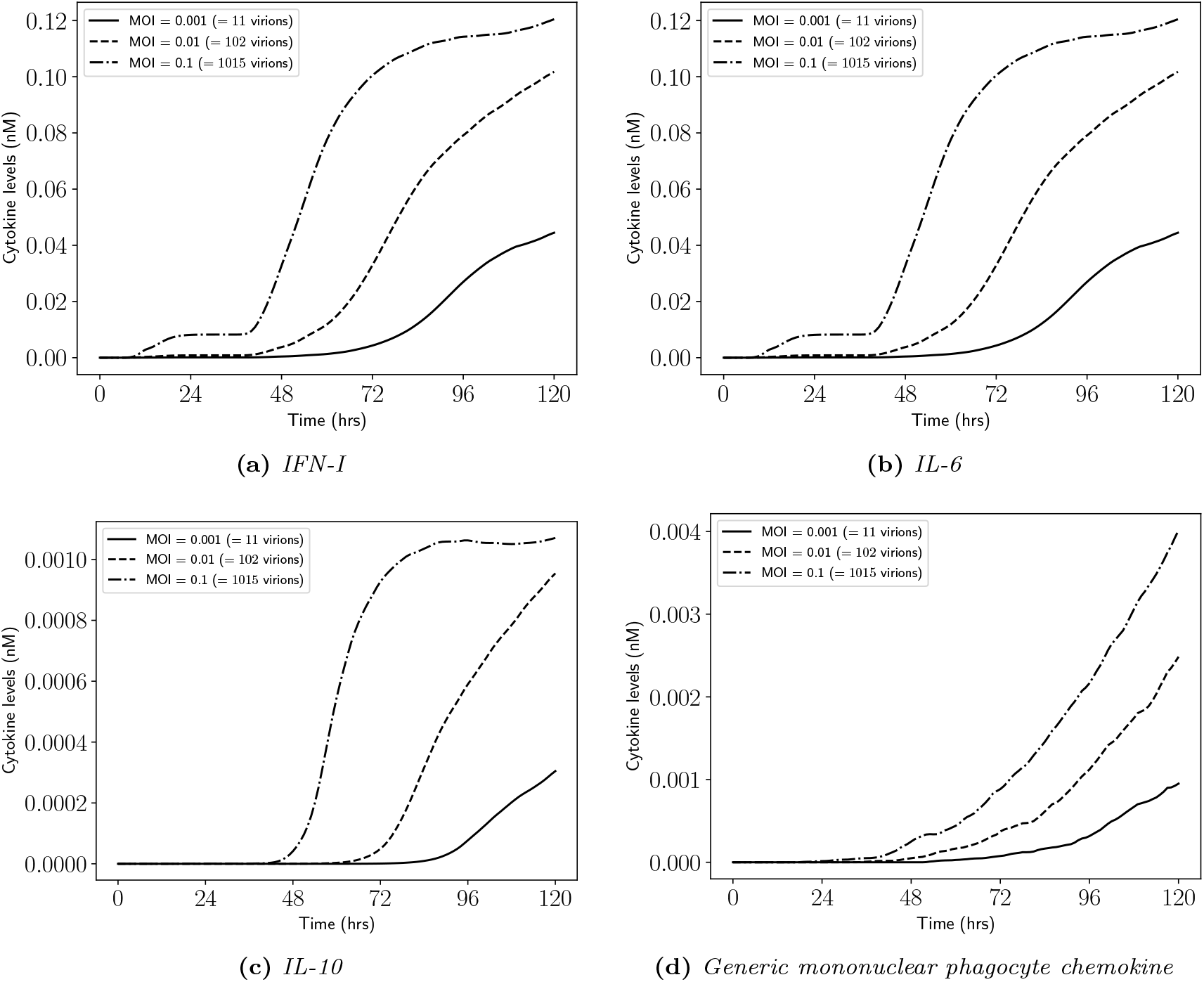
(Cytokine levels for increasing MOI) *Illustration of (a) type I interferon, (b) interleukin 6, (c) interleukin 10 and (d) generic mononuclear phagocyte chemokine levels as the simulation progresses, for increasing MOI. Baseline parameters are given in Tables 1-4*.

Figure 7 illustrates the intracellular and extracellular viral load per grid, as the simulation progresses, for each MOI considered. In Fig. 7(a) a local maximum can be observed for all MOI values; as the MOI is increased, we see the local maximum appear earlier in the simulation, but the magnitude of the local maximum does not monotonically increase as the MOI is increased. Indeed, for MOI = 0.01, we see a greater local maximum than when MOI = 0.1. Our hypothesis is that the local maximums appear due to a sudden increase in IFN-I and are therefore, affected by both the delay in epithelial IFN-I secretion as well as the epithelial IFN-I secretion rate, and is discussed in more detail in §3.2. It is interesting, therefore, that the local maximum may be overcome to produce a rebound in intracellular virions per grid, where the higher the MOI, the steeper the rebound, and may be caused by viral replication inside epithelial cells where the local extracellular IFN-I levels are lower, or because the IFN-I inhibition of viral entry and repliation has saturated whilst the local extracellular viral load is increasing. In Fig. 7(b), it appears that for *t* < 48 hours, the extracellular viral loads are approximately similar for each MOI considered. However, for *t* > 48 hours, the viral loads begin to differ (although not substantially until *t* = 96 hours) and by the end of the simulation we observe an increasing number of extracellular virions as the MOI is increased, as expected. Around *t* = 96 hours, for MOI = 0.001, we see a temporary flattening of the number of extracellular virions which correlates with the local maximum in intracellular virions per grid (Fig. 7(a)).

Figure 8 illustrates the levels of each cytokine considered as the simulation progresses, for each MOI value. In all cases, we see an increasing amount of cytokine by the end of the simulation as the MOI is increased; similarly, the initial increase in cytokine level occurs earlier in the simulation, and generally has a steeper increase, as the MOI is increased. For MOI = 0.1, we see a sharp increase IFN-I levels around *t* = 48 hours, which correlates with the observed local maximum in the intracellular viral load per grid (Fig. 7(a)). However, we do not observe any changes in cytokine levels that correspond to the rebound in intracellular viral load per grid. IL-10 levels (Fig. 8(c)) seem to flatten off towards the end of the simulation for the largest MOI considered, due to a short flattening in IL-6 levels (Fig. 8(b)), whilst are still increasing for the other MOI values.

### 3.2 Delayed IFN-I secretion from epithelial cells

A typical observation following coronavirus infection of epithelial cells is a delayed type 1 interferon (IFN-I) response [44, 45], which has been suggested to increase disease severity [42, 21]. Therefore, we investigate the impact of a delayed IFN-I secretion from epithelial cells on the spread of infection, epithelial and macrophage numbers, as well as viral load and cytokine levels. We consider four values of the epithelial IFN-I secretion delay (*S*_50,*V*_ = 1,10,100,1000 virions) so that IFN-I is only secreted from infected and infectious epithelial cells when 1, 10, 100 or 1000 virions are detected in the cell cytoplasm. Note that we fix the MOI = 0.01 in this section, and do not consider delays in IFN-I inhibition of viral entry and replication pathways.

Figure 9 illustrates the spread of infection at the end of the simulation (*T* = 120 hours), for fixed MOI = 0.01, as the epithelial IFN-I secretion delay is increased. Clearly, we observe an increase in the spread of infection for longer epithelial IFN-I secretion delays, as the viral entry and replication processes are able to proceed for longer without being inhibited by IFN-I. Similar to Fig. 5, we observe mostly active and apoptotic macrophages at the site of infection, with resting macrophages mostly localised to the boundary of the infection and away from it. As the epithelial IFN-I secretion delay is increased, we see an increasing number of resting and active macrophages but a decrease in apoptotic macrophages due to lower levels of IFN-I.

**Figure 9:**
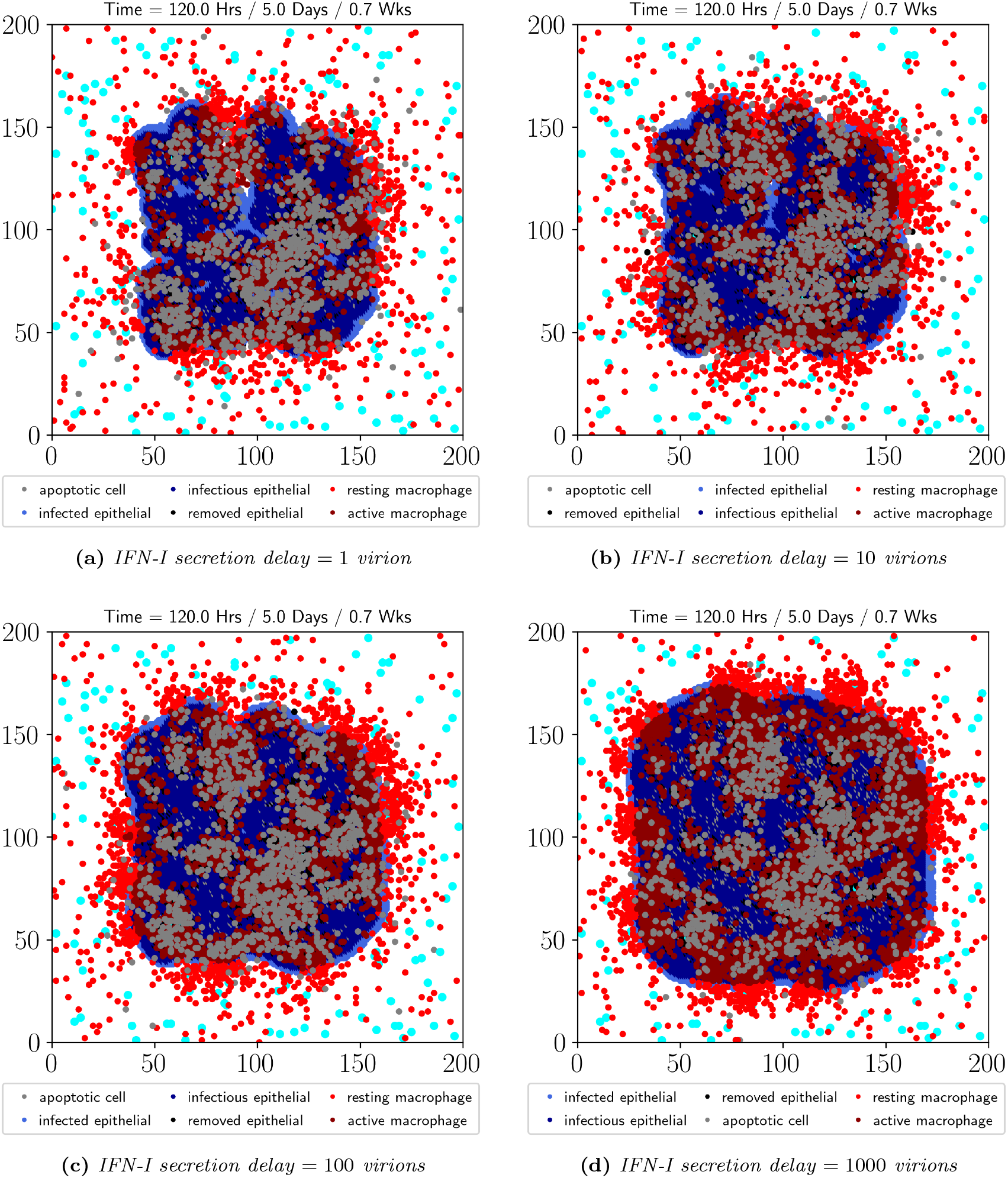
(Spread of infection for increasing epithelial IFN-I secretion delay) *Illustration of the spread of infection at the end of the simulation (T* = 120 *hours) for epithelial IFN-I secretion delay (a) S*_50,*V*_ = 1 *virion, (b) S*_50,*V*_ = 10 *virions, (c)* S_50,*V*_ = 100 *virions, and (d) S*_50,*V*_ = 1000 *virions. Note that the MOI* = 0.01 *and the other baseline parameter values are given in Tables 1-4*.

Figures 10(a)-(c) illustrate the number of epithelial cells in an infectious, apoptotic or removed state, respectively, for each value of the epithelial IFN-I secretion delay. As expected, in Fig. 10(a), we see an increasing number of infectious epithelial cells by the end of the simulation, for increasing IFN-I secretion delay due to the enhanced spread illustrated in Fig. 9. Additionally, for *t* < 48 hours, we do not observe significant differences between the number of infectious, apoptotic or removed epithelial cells, for each IFN-I secretion delay. Epithelial cells may be placed into an apoptotic state due to high intracellular viral loads or due to high IFN-I; in Fig. 10(b) we see an increase in apoptotic epithelial cells at the end of the simulation, as the IFN-I secretion delay is increased, due to an increase in intracellular viral load (see Fig. 11(a)). Epithelial cells may be placed into a removed state after successful macrophage phagocytosis; in Fig. 10(c) we see an increase in removed epithelial cells at the end of the simulation, as the IFNI secretion delay increases, which correlates with an increase in active macrophage numbers (see Fig. 10(e)) for each IFN-I secretion delay considered. Interestingly, the number of resting macrophages seems to approximately flatten after *t* > 72 hours (Fig. 10(d)). Furthermore, reduced apoptotic macrophages are seen for the longest epithelial IFN-I secretion delay (Fig. 10(f)), as illustrated in Fig. 9.

**Figure 10:**
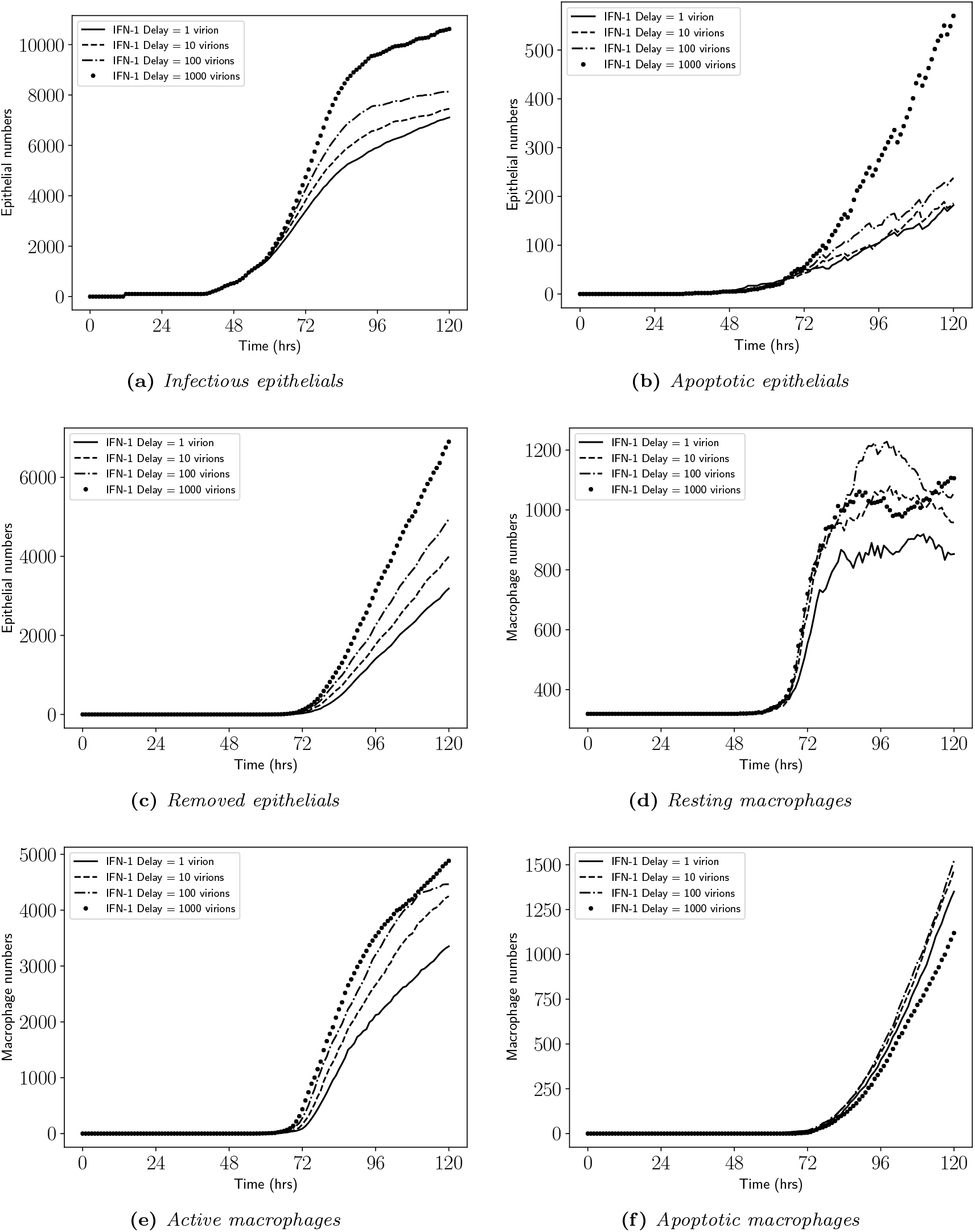
(Cell numbers for increasing epithelial IFN-I secretion delay) *Number of (a) infectious, (b) apoptotic and (c) removed epithelial cells, as well as (d) resting, (e) active and (f) apoptotic macrophage numbers, as the simulation progresses, for increasing IFN-I secretion delay* (*S*_50,*V*_). *Note that the MOI* = 0.01 *and the other baseline parameters are given in Tables 1-4*.

**Figure 11:**
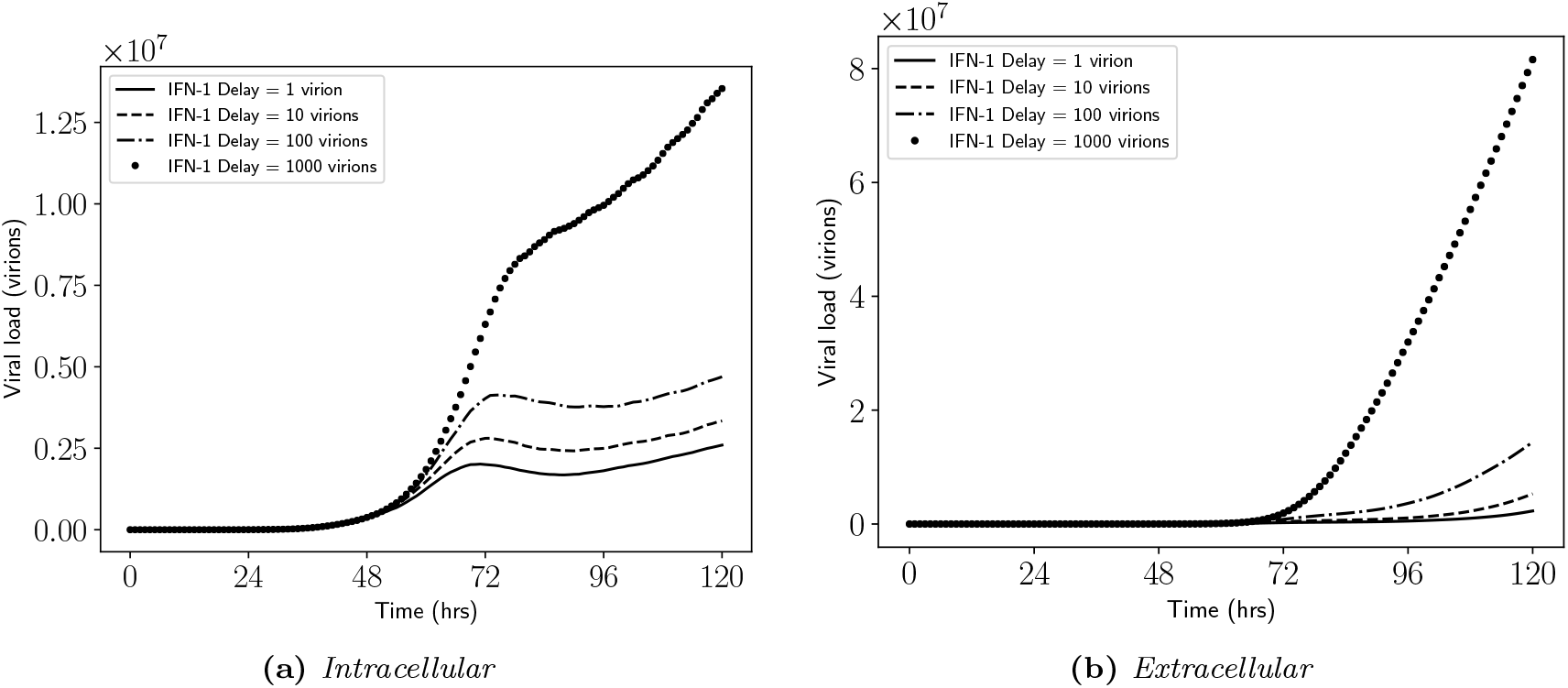
(Viral loads per grid for increasing epithelial IFN-I secretion delay) *Illustration of (a) intracellular and (b) extracellular viral loads per grid as the simulation progresses, for increasing IFN-I secretion delay* (*S*_50,*V*_). *Note that the MOI* = 0.01 *and the other baseline parameters are given in Tables 1-4*.

Figure 11 illustrates the intracellular and extracellular viral load per grid as the simulation progresses, for each IFN-I secretion delay considered. For both the intracellular (Fig. 11(a)) and extracellular (Fig. 11(b)) viral loads per grid, we see a very significant increase by the end of the simulation, for the longest IFN-I secretion delay (= 1000 virions). Additionally, the local maximum that can be observed in the intracellular viral load per grid for epithelial IFN-I secretion delays of 1, 10, 100 virions, cannot be observed for an IFN-I secretion delay of 1000 virions; instead, we observe a decrease in the growth rate of intracellular viral load per grid, further suggesting that IFN-I (relative to the amount of intracellular virus) is responsible for the local maximum in the intracellular viral load per grid observed earlier (Fig. 7(a)). Therefore, the rebound in intracellular viral load per grid observed earlier (Fig. 7(a)), is still observed for IFN-I secretion delays of 1,10,100 virions, but not for a IFN-I secretion delay of 1000 virions (Fig. 11(a)).

Figure 12 illustrates extracellular IFN-I, IL-6, IL-10 and generic mononuclear phagocyte chemokine levels as the simulation progresses, for each epithelial IFN-I secretion delay considered. As expected, IFN-I levels decrease as the secretion delay increases (Fig. 12(a)), but IL-6 levels increase as IFN-I secretion delay increases (Fig. 12(b)) due to the increasing spread of infection and thus, infectious epithelial cells (Figs. 9 and 10(a)). Correspondingly, we see an increase in IL-10 levels as IFN-I secretion delay increases (Fig. 12(c)) due to the increase in IL-6 levels; however, for the longest IFN-I secretion delay, IL-10 levels seem to be declining towards the end of the simulation. Additionally, we observe an increase in generic mononuclear phagocyte chemokine as IFN-I secretion delay increases (Fig. 12(d)) due to an increase in the total number of apoptotic epithelial cells and macrophages (Figs. 10(b),(f)).

**Figure 12:**
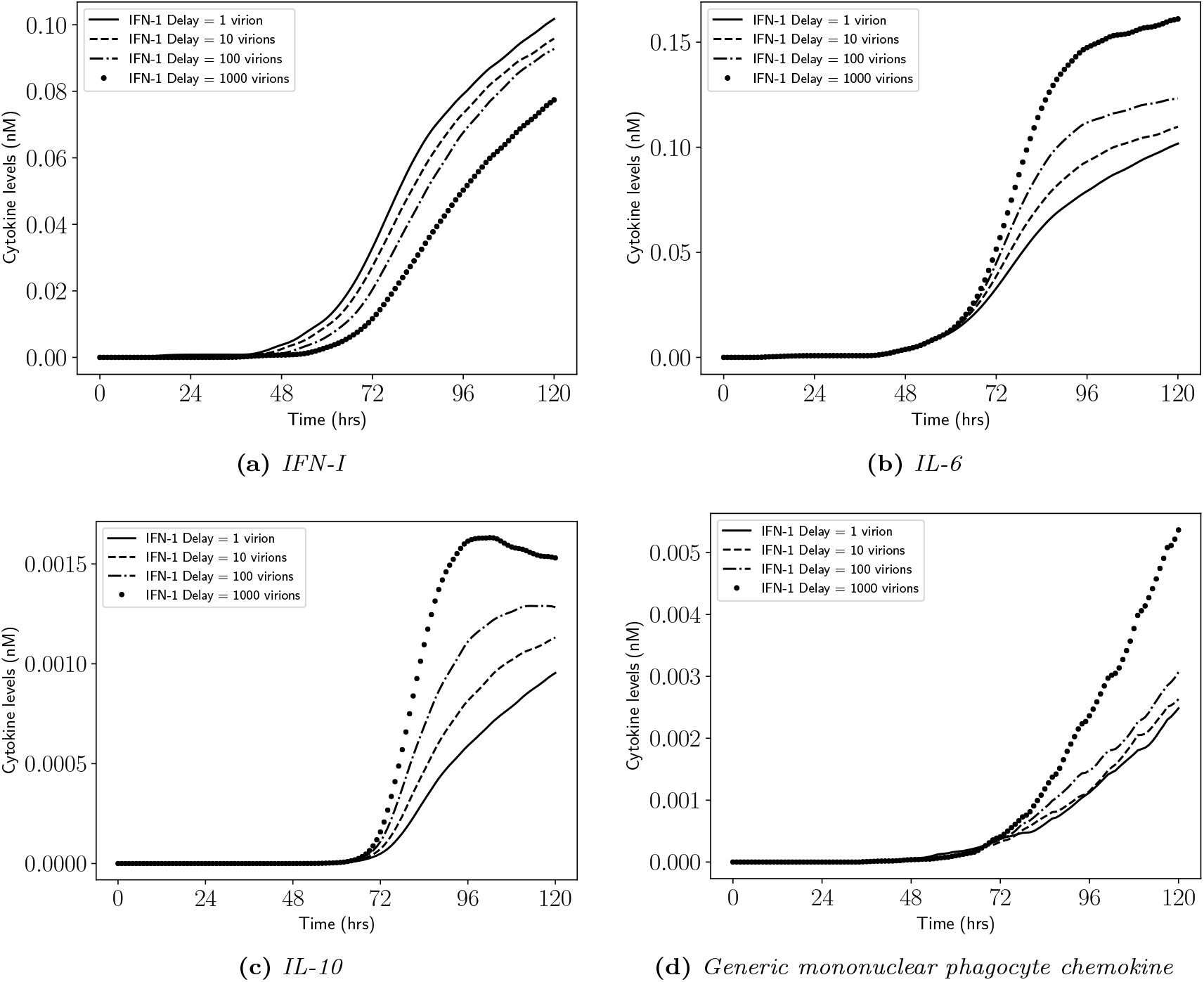
(Cytokine levels for increasing epithelial IFN-I secretion delay) *Illustration of (a) type I interferon, (b) interleukin 6, (c) interleukin 10 and (d) generic mononuclear phagocyte chemokine levels as the simulation progresses, for increasing IFN-I secretion delay* (*S*_50,*V*_). *Note that the MOI* = 0.01 *and the other baseline parameters are given in Tables 1-4*.

To further investigate the influence of IFN-I secretion on the spread of infection, epithelial and macrophage numbers, as well as viral load per grid and cytokine levels, we consider three values of the epithelial IFN-I secretion rate 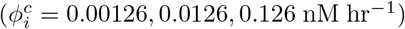, for fixed MOI = 0.01 and epithelial IFN-I secretion delay = 1 virion. Figure 13 illustrates the spread of infection at the end of the simulation (*T* = 120 hours), as the epithelial IFN-I secretion rate increases. Clearly, as the epithelial IFN-I secretion rate is increased, we see a decreased number of infectious epithelial cells, as well as a decrease in resting, active and apoptotic macrophages at the site of infection; however, there is an increase in macrophage apoptosis away from the site of infection (Fig. 13(c)) due to larger amounts of extracellular IFN-I.

**Figure 13:**
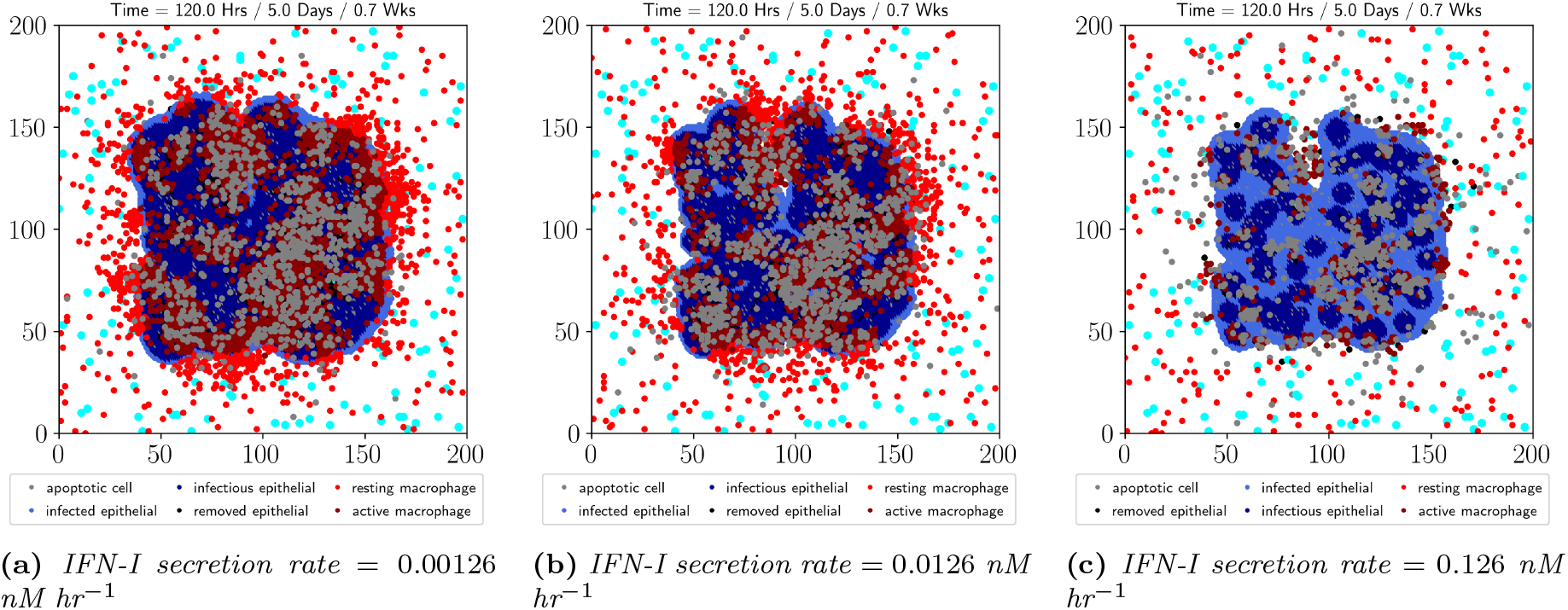
(Spread of infection for increasing epithelial IFN-I secretion rate) *Illustration of the spread of infection at the end of the simulation (T* = 120 *hours) for epithelial IFN-I secretion rate (a)* 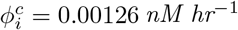, *(b)* 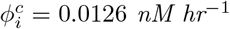, *and (c)* 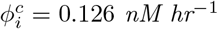. *Note that the MOI* = 0.01, *epithelial IFN-I secretion delay* = 1 *virion, and the other baseline parameter values are given in Tables 1-4*.

Figures 14(a),(b),(c) illustrate the infectious, apoptotic and removed epithelial cell numbers, as the simulation progresses, for each IFN-I secretion rate considered. As shown in Fig. 13, we see a decreasing number of infectious epithelial cells, at the end of the simulation, as IFN-I secretion rate increases (Fig. 14(a)). However, for *t* < 48 hours we do not see any significant differences in the number of infectious, apoptotic or removed epithelial cells, for each IFN-I secretion rate considered. As epithelial cells may be placed into an apoptotic state because of high IFN-I or high intracellular virus, we observe higher numbers of apoptotic epithelial cells for the smallest and largest IFN-I secretion rates considered, when compared to the median IFN-I secretion rate; however, for the largest IFN-I secretion, we see a significantly higher number of apoptotic epithelial cells (Fig. 14(b)), by the end of the simulation. In Fig. 14(c), we see a decreasing number of removed epithelial cells at the end of the simulation, as the IFN-I secretion rate is increased, which correlates with the decreasing levels of active macrophages (Fig. 14(e)). For the largest IFN-I secretion rate, we do not observe an expansion of resting macrophages (Fig. 14(d)) due to the lower levels of IL-6 (Fig. 16(b)); therefore, for *t* > 48 hours, we see a decrease in the number of resting macrophages as the simulation progresses due to activation and apoptosis. For the other IFN-I secretion rates, we observe an expansion of resting macrophages with the greatest expansion observed for the smallest IFN-I secretion rate. Due to the lower levels of macrophages observed for the highest IFN-I secretion rate, we observe lower levels of apoptotic macrophages (Fig. 14(f)); however, for the other IFN-I secretion rates considered, we do not see any difference in apoptotic macrophage numbers over the course of the simulation.

**Figure 14:**
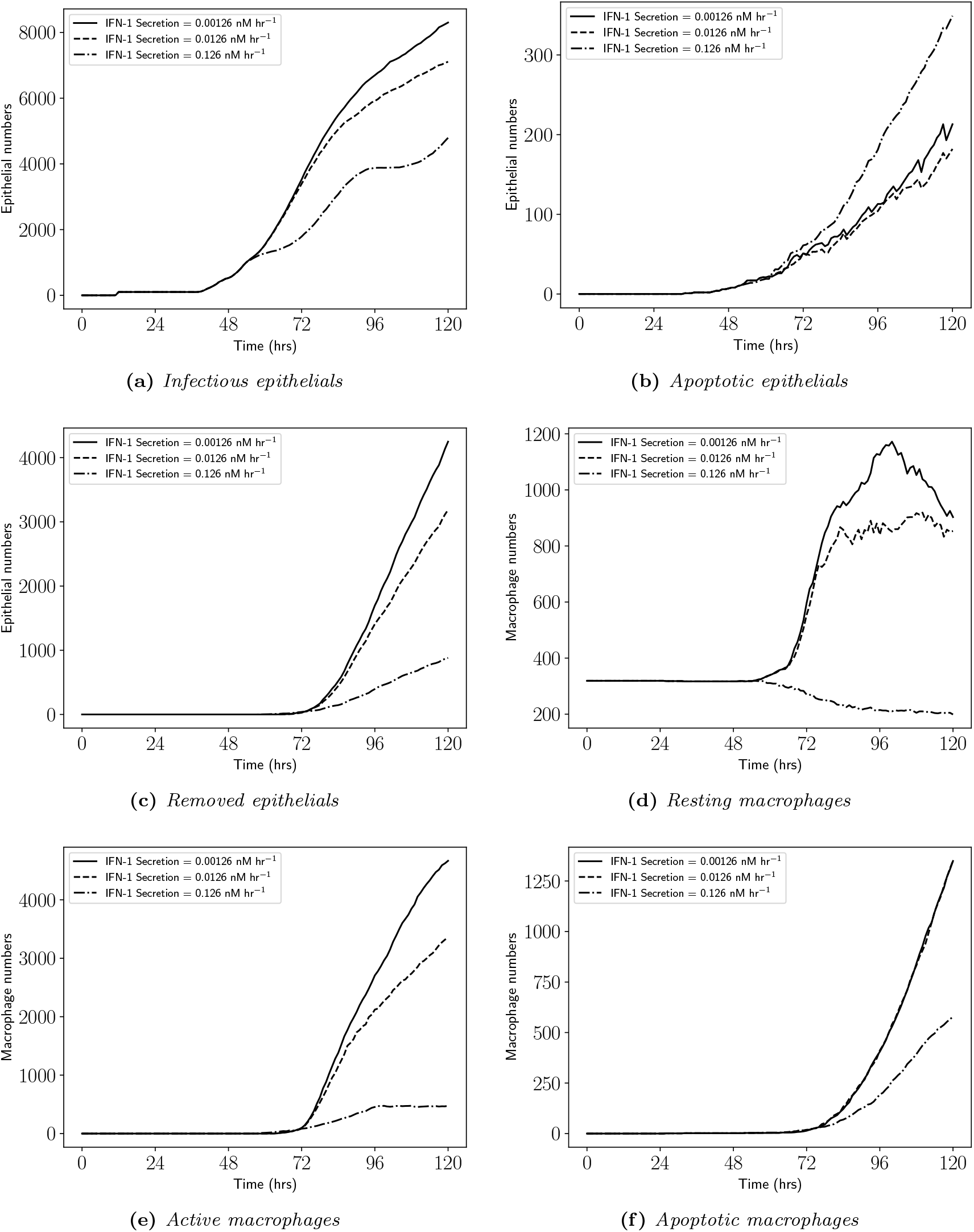
(Cell numbers for increasing epithelial IFN-I secretion rate) *Number of (a) infectious, (b) apoptotic and (c) removed epithelial cells, as well as (d) resting, (e) active and (f) apoptotic macrophage numbers, as the simulation progresses, for increasing epithelial IFN-I secretion rate* 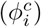. *Note that the MOI* = 0.01, *epithelial IFN-I secretion delay* = 1 *virion, and the other baseline parameters are given in Tables 1-4*.

Figure 15 illustrates the intracellular and extracellular viral loads per grid, as the simulation progresses, for each epithelial IFN-I secretion rate considered. In Fig. 15(a), we see an increasing intracellular viral load per grid with increasing IFN-I secretion rate, by the end of the simulation. Interestingly, for the smallest IFN-I secretion rate (0.00126 nM hr^-1^) we do not observe a local maximum in intracellular viral load, which can be seen for the other IFN-I secretion rate values; instead, similar to the longest epithelial IFN-I secretion delay (Fig. 11(a)), we observe a decrease in the growth rate of intracellular viral load per grid. Additionally, following the local maximum, the rebound in the intracellular viral load which has been observed previously (e.g. Fig. 7(a)), is not clearly observed for the largest IFN-I secretion value.

**Figure 15:**
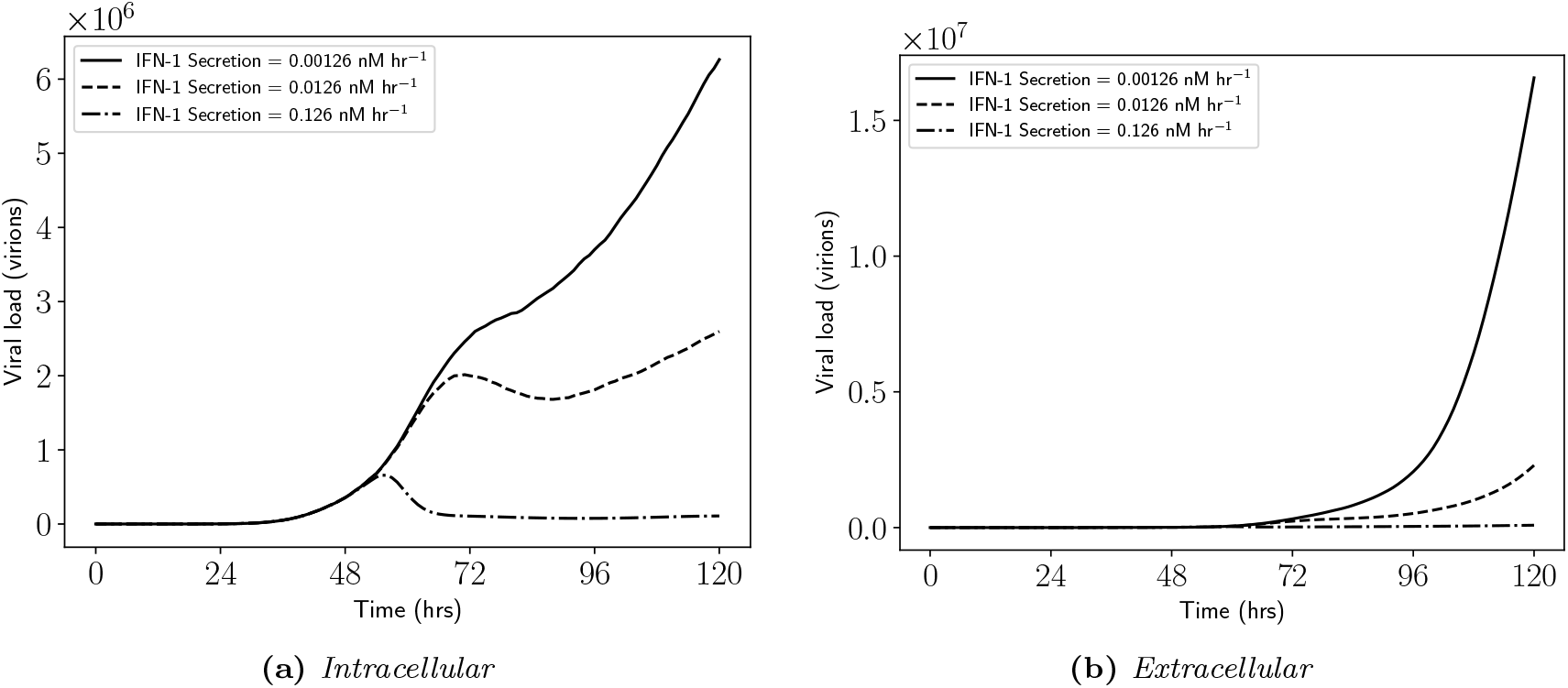
(Viral loads per grid for increasing epithelial IFN-I secretion rate) *Illustration of (a) intracellular and (b) extracellular viral loads per grid as the simulation progresses, for increasing IFN-I secretion rate* 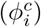. *Note that the MOI* = 0.01, *epithelial IFN-I secretion delay* = 1 *virion, and the other baseline parameters are given in Tables 1-4*.

Figure 16 illustrates the IFN-I, IL-6, IL-10 and generic mononuclear phagocyte chemokine levels, as the simulation progresses, for each epithelial IFN-I secretion rate considered. As expected, increasing the IFN-I secretion rate increases the IFN-I levels (Fig. 16(a)); higher IFN-I levels reduces the number of infectious epithelial cells (Fig. 14(a)), resulting in decreasing IL-6 and IL-10 levels as the IFN-I secretion rate is increased (Figs. 16(b),(c)). We observe slightly higher levels of generic mononuclear phagocyte chemokine for the largest IFN-I secretion rate (Fig. 16(d)), which correlates with a higher total number of apoptotic cells (Fig. 14(b),(f)); however, we do not correspondingly observe an increase in macrophage numbers for increasing IFN-I secretion rates (Fig. 14(d)), even though the generic mononuclear phagocyte chemokine is involved in the recruitment of macrophages, due to insufficient IL-6 levels.

**Figure 16:**
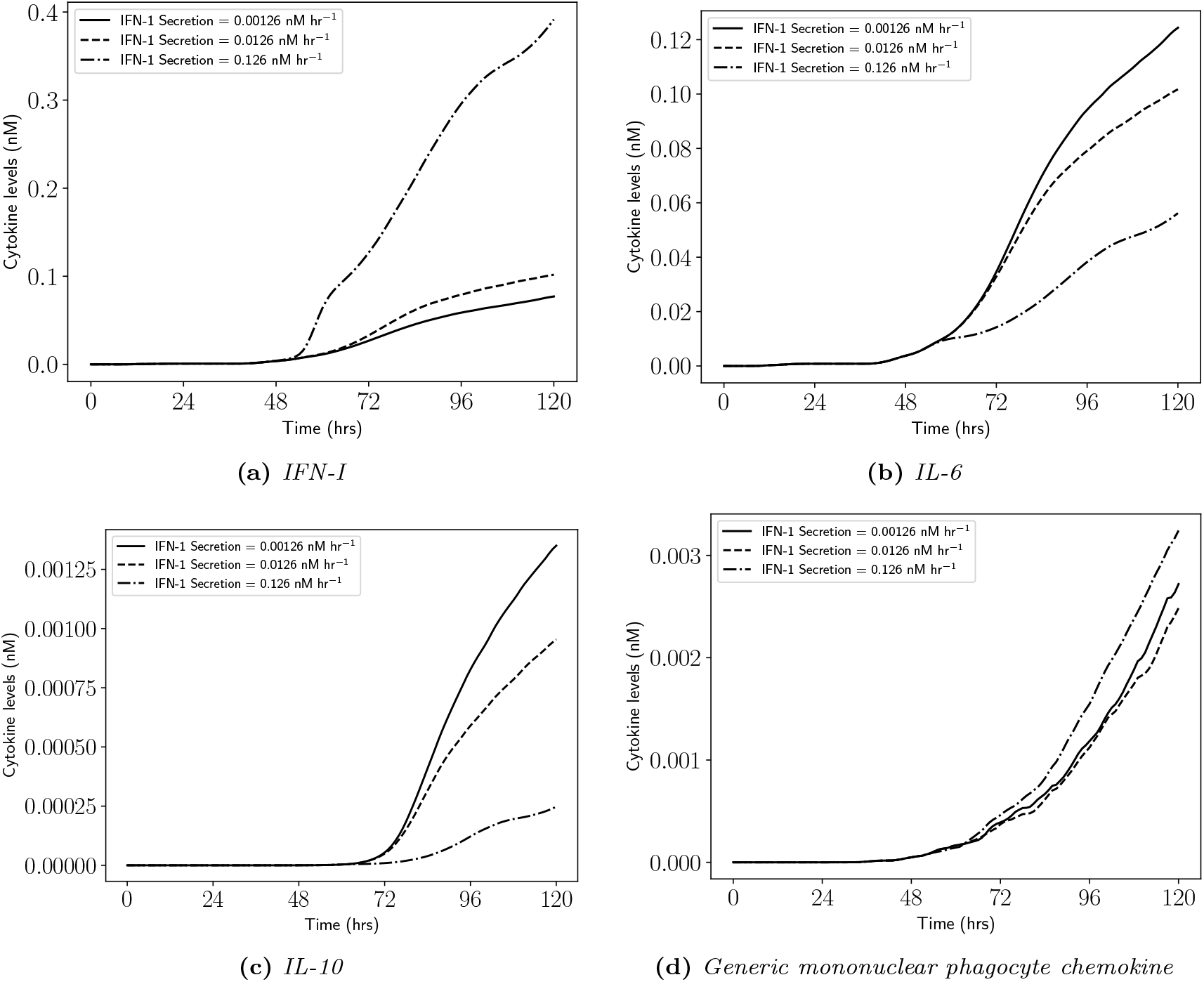
(Cytokine levels for increasing epithelial IFN-I secretion rate) *Illustration of (a) type I interferon, (b) interleukin 6, (c) interleukin 10 and (d) generic mononuclear phagocyte chemokine levels as the simulation progresses, for increasing IFN-I secretion rate* 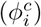. *Note that the MOI* = 0.01, *epithelial IFN-I secretion delay* = 1 *virion, and the other baseline parameters are given in Tables 1-4*.

### 3.3 Macrophage parameter variations

We consider variations of two macrophage parameters in our simulation: macrophage virus internalisation rate and macrophage activation half-max. Figure 17 illustrates the spread of infection at the end of the simulation, as the macrophage virus internalisation rate increases. There does not appear to be any significant differences between the spread of infection at the end of the simulation. This is confirmed in Figs. 18(a),(b),(c), where we see similar amounts infectious, apoptotic and removed epithelial cells, by the end of the simulation, as the macrophage virus internalisation rate increases. However, in Fig. 17, we do observe (slight) decreases in macrophage numbers as the virus internalisation rate increases. This is confirmed in Figs. 18(d),(e),(f), where we observe decreasing levels of resting, active and apoptotic macrophages as the virus internalisation rate increases. Additionally, we observe only slight differences in intracellular and extracellular viral loads per grid (Fig. 19), as well as only slight differences in the cytokine levels (Fig. 20).

**Figure 17:**
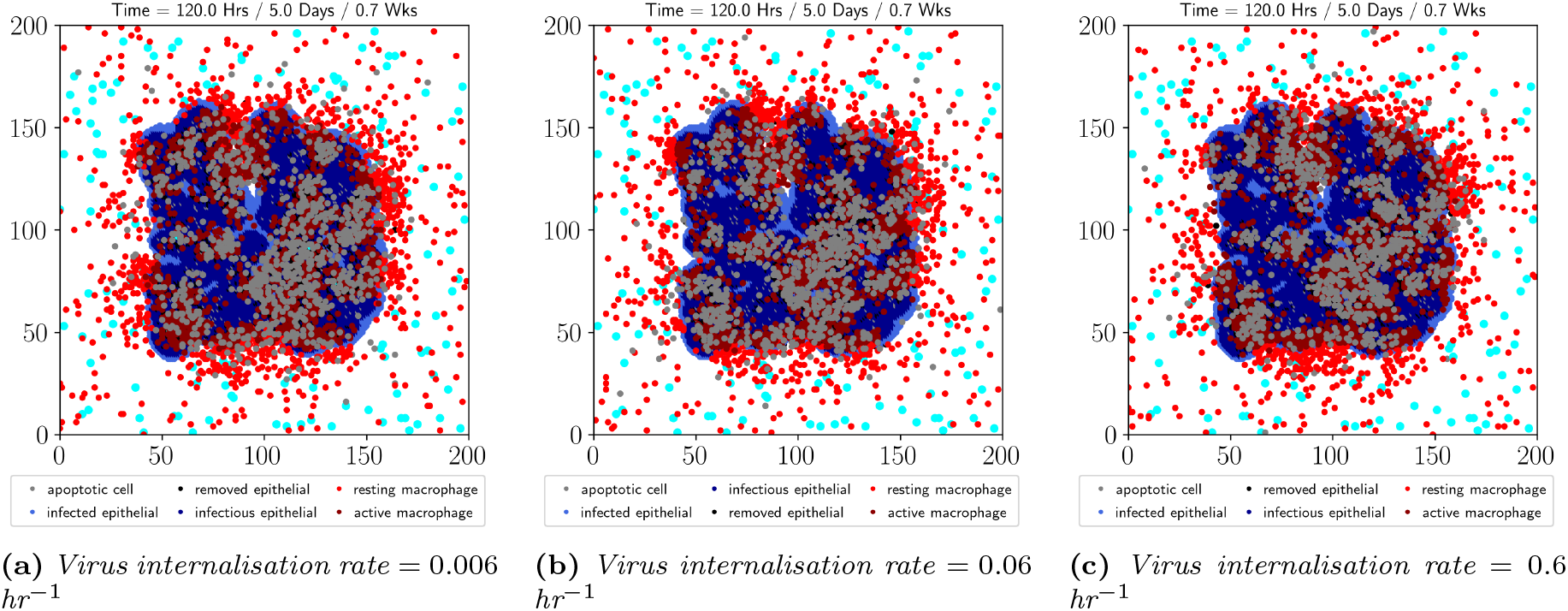
(Spread of infection for increasing macrophage virus internalisation rate) *Illustration of the spread of infection at the end of the simulation (T* = 120 *hours) for macrophage virus internalisation rate (a) r*_[*V*]_ = 0.006 *hr*^−1^, *(b) r*_[*V*]_ = 0.06 *hr*^−1^, *and (c) r*_[*V*]_ = 0.6 *hr*^−1^. *Note that the MOI* = 0.01 *and the other baseline parameter values are given in Tables 1-4*.

**Figure 18:**
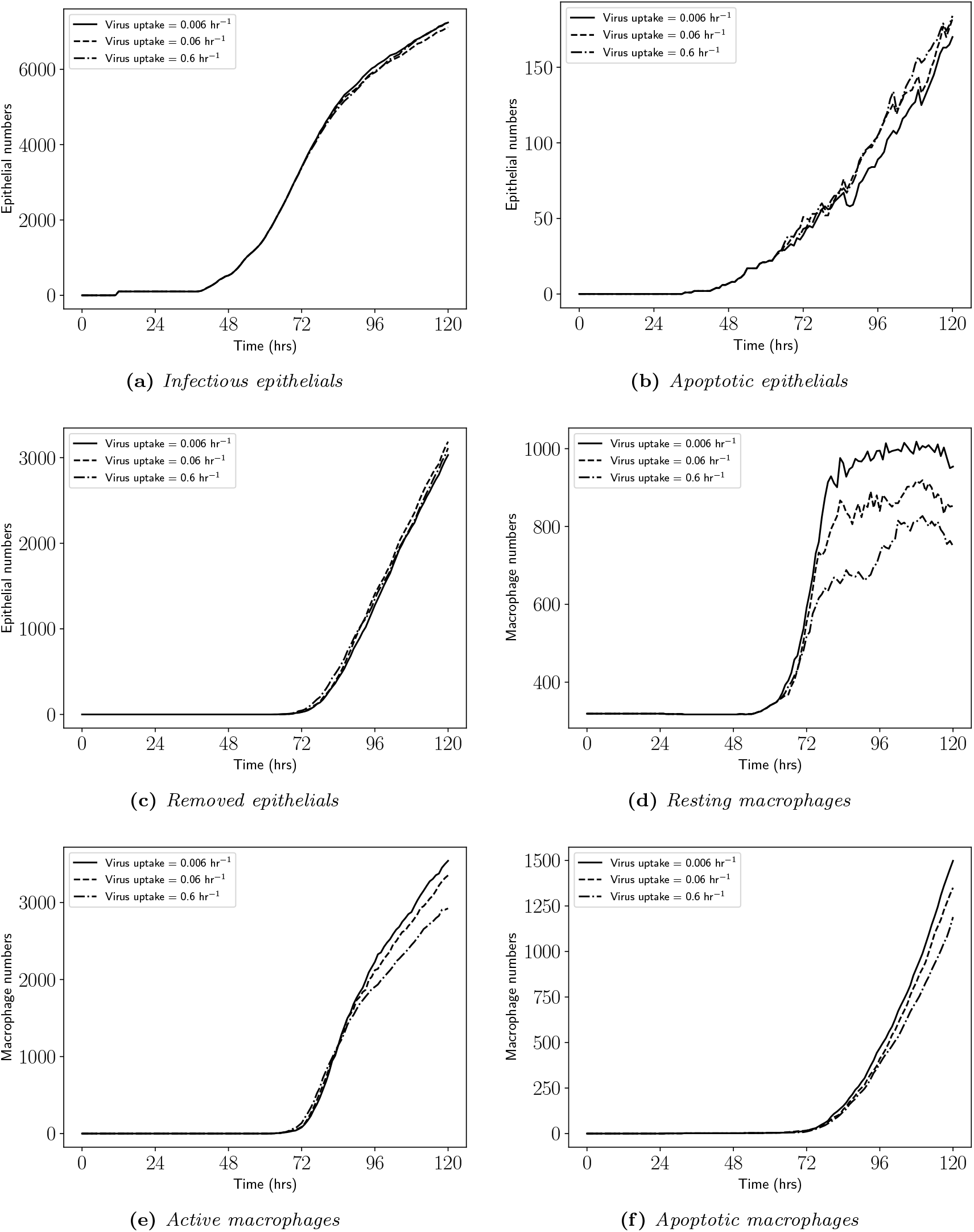
(Cell numbers for increasing macrophage virus internalisation rate) *Number of (a) infectious, (b) apoptotic and (c) removed epithelial cells, as well as (d) resting, (e) active and (f) apoptotic macrophage numbers, as the simulation progresses, for increasing macrophage virus internalisation rate* (*r*_[*V*]_). *Note that the MOI* = 0.01 *and the other baseline parameters are given in Tables 1-4*.

**Figure 19:**
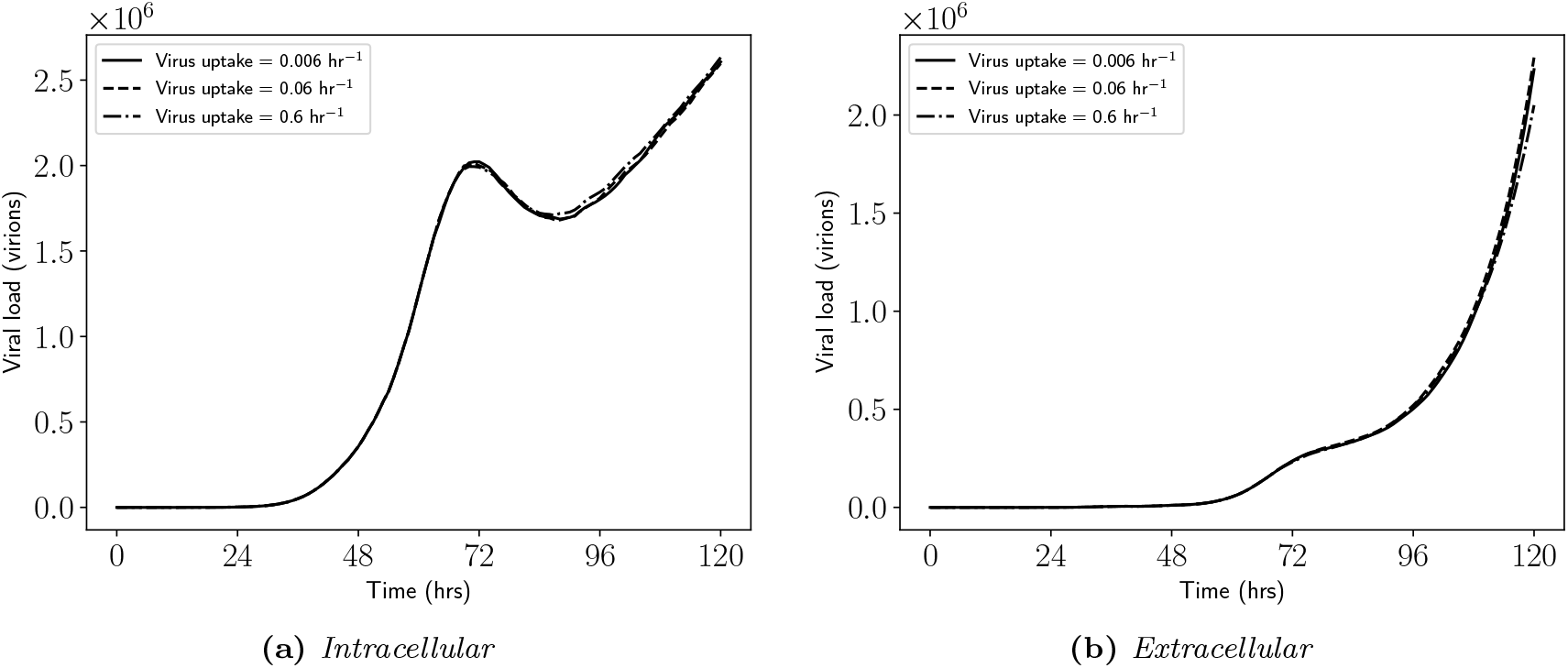
(Viral loads per grid for increasing macrophage virus internalisation rate) *Illustration of (a) intra-cellular and (b) extracellular viral loads per grid, as the simulation progresses, for increasing macrophage virus internalisation rate* (*r*_[*V*]_). *Note that the MOI* = 0.01 *and the other baseline parameters are given in Tables 1-4*.

**Figure 20:**
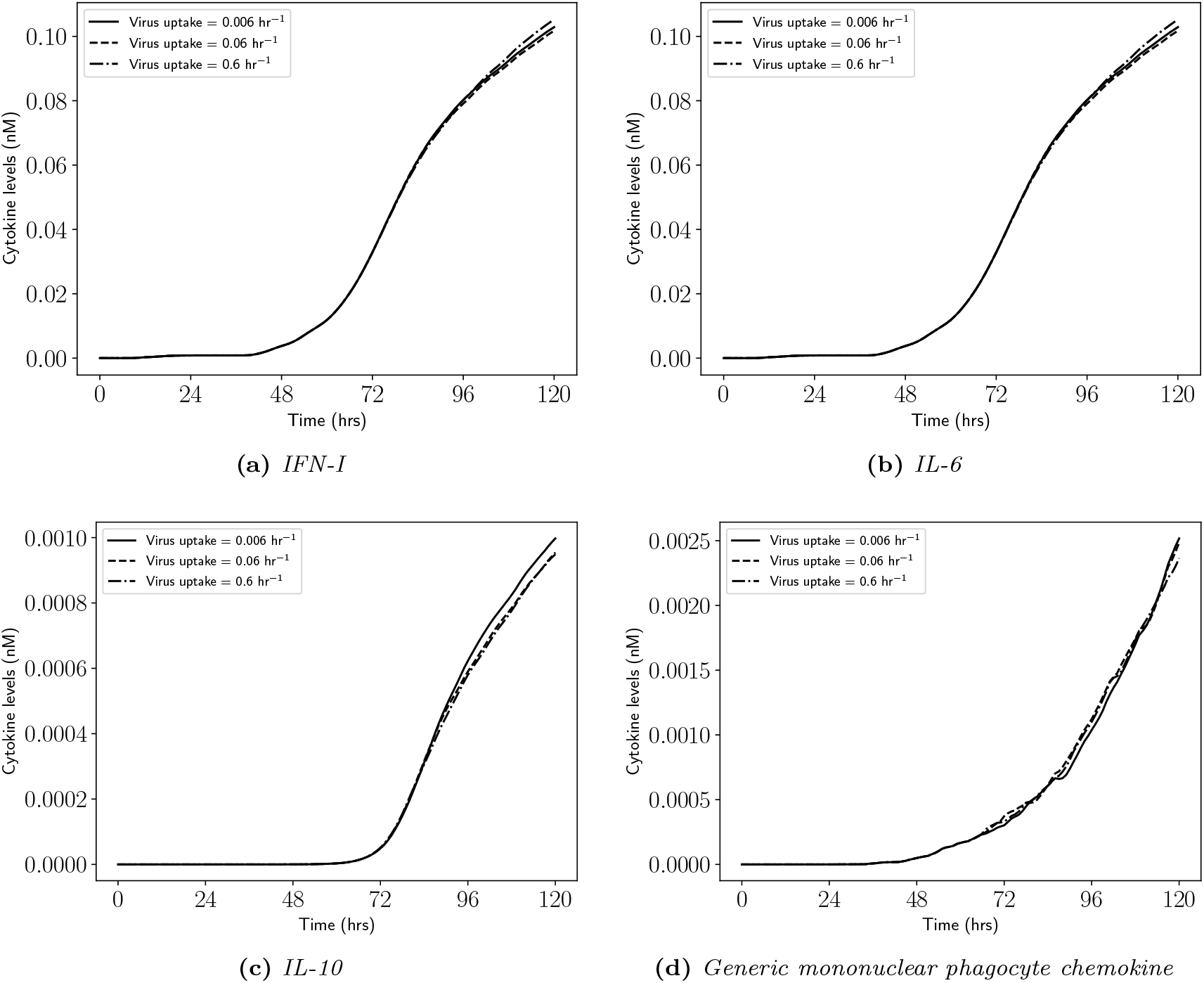
(Cytokine levels for increasing macrophage virus internalisation rate) *Illustration of (a) type I interferon, (b) interleukin 6, (c) interleukin 10 and (d) generic mononuclear phagocyte chemokine levels as the simulation progresses, for increasing macrophage virus internalisation rate* (*r*_[*V*]_). *Note that the MOI* = 0.01 *and the other baseline parameters are given in Tables 1-4*.

Figure 21 illustrates the spread of infection at the end of the simulation, as the macrophage activation half-max increases 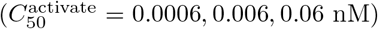. The spread of infection (measured by infectious epithelial cells) appears similar for all activation half-max values, however as the activation half-max value is increased we see a significant increase in the number of resting and apoptotic macrophages. Furthermore, a greater proportion of active macrophages may be found away from the site of infection for the lowest activation half-max value.

**Figure 21:**
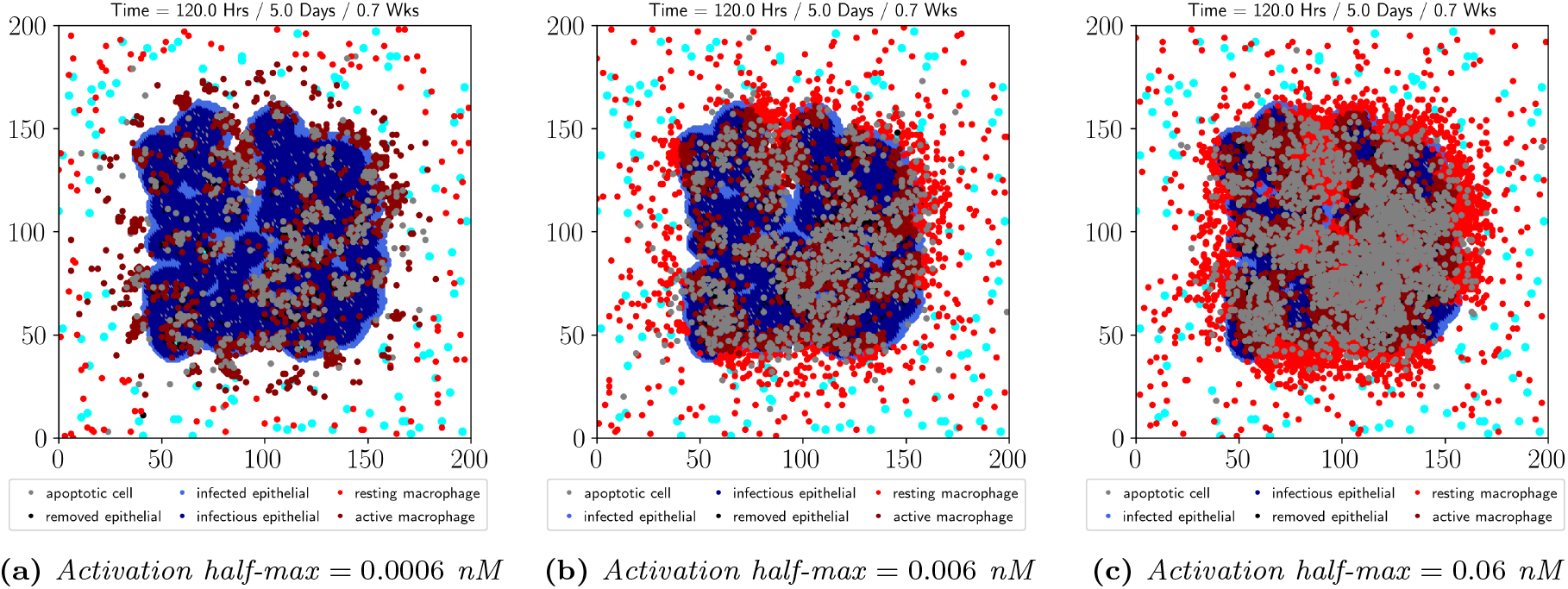
(Spread of infection for increasing macrophage activation half-max) *Illustration of the spread of infection at the end of the simulation (T* = 120 *hours) for macrophage activation half-max (a)* 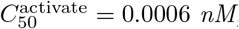 *nM, (b)* 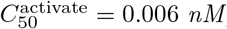 *nM, and (c)* 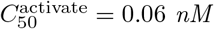 *nM. Note that the MOI* =0.01 *and the other baseline parameter values are given in Tables 1-4*.

Figure 22 illustrates the number of infectious, apoptotic and removed epithelial cells, as well as resting, active and apoptotic macrophage numbers, as the simulation progresses, for increasing activation half-max values. For the lowest activation half-max value (meaning macrophages can activate readily), we observe higher numbers of infectious and apoptotic epithelial cells (Figs. 22(a),(b)); however, we see a lower level of removed epithelial cells (Fig. 22(c)) due to lower numbers of active macrophages (Fig. 22(e)). For the lowest activation half-max value, there does not appear to be a significant expansion of macrophages (Figs. 22(d),(e)). Interestingly, for the highest activation half-max value considered, there appears to be a flattening of infectious and apoptotic epithelial cells (Figs. 22(a),(b)); however, we clearly see a larger number of removed epithelials (Fig. 22(c)) due to higher levels of active macrophages (Fig. 22(e)), which are most likely observed due to the higher levels of resting macrophages (Fig. 22(d)).

**Figure 22:**
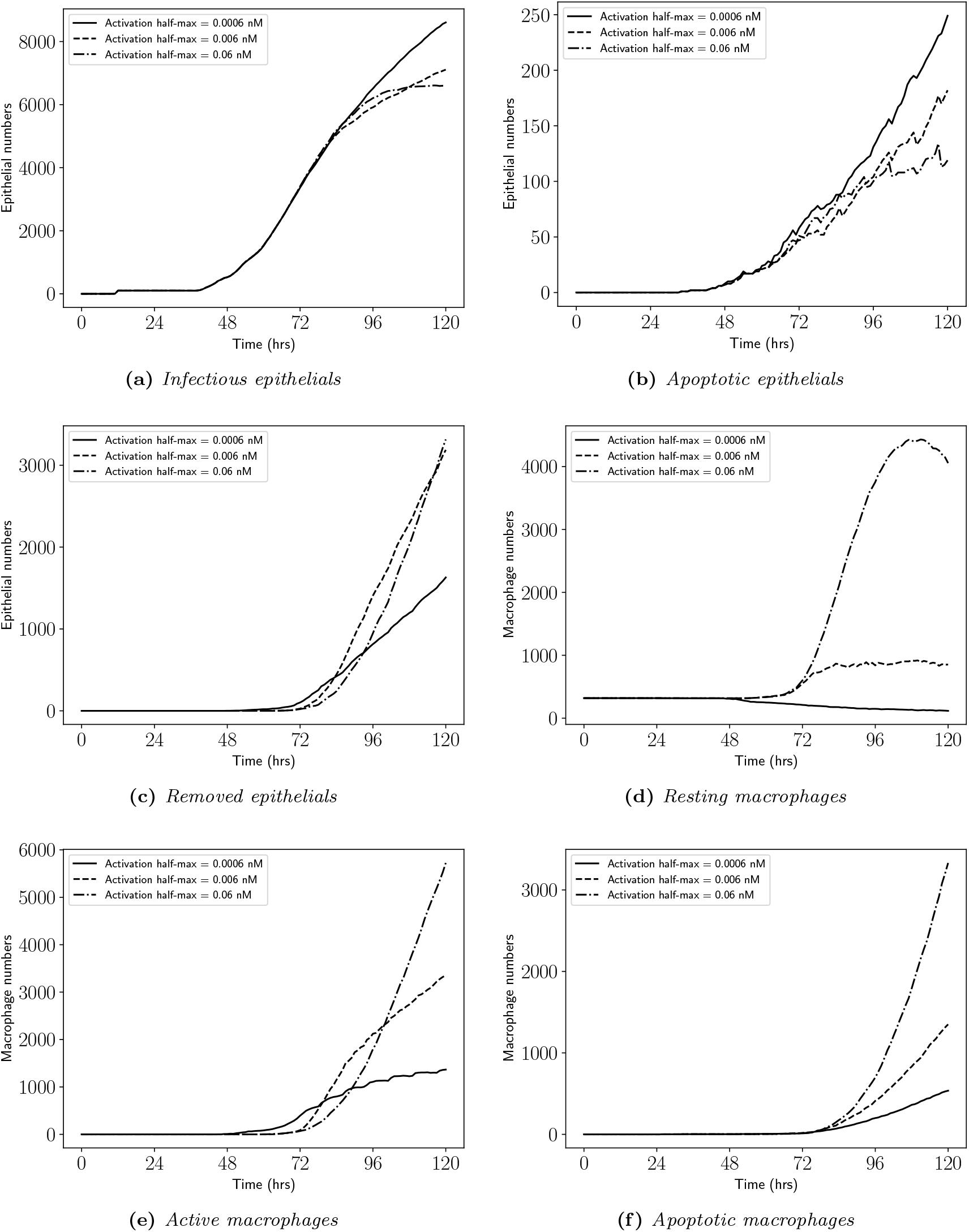
(Cell numbers for increasing macrophage activation half-max) *Number of (a) infectious, (b) apoptotic and (c) removed epithelial cells, as well as (d) resting, (e) active and (f) apoptotic macrophage numbers, as the simulation progresses, for increasing macrophage activation half-max 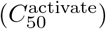. Note that the MOI* = 0.01 *and the other baseline parameters are given in Tables 1-4*.

Figure 23 illustrates the intracellular and extracellular viral loads per grid, as the simulation progresses, for all activation half-max values considered. The intracellular viral loads per grid follow similar trajectories up to ≈ *t* = 72 hours, for each activation half-max value considered; however, for *t* > 72 hours, the viral load per grid for the largest activation half-max value reaches a lower minimum than for the other activation half-max values (Fig. 23(a)). However, there does not appear to be any significant changes in the IFN-I levels around *t* = 72 hours that would explain the lower minimum observed in the intracellular viral load for the largest activation half-max value. Similarly, we observe a lower extracellular viral load per grid for the largest activation half-max value (Fig. 23(b)), due to the larger number of active macrophages (Fig. 22(e)).

**Figure 23:**
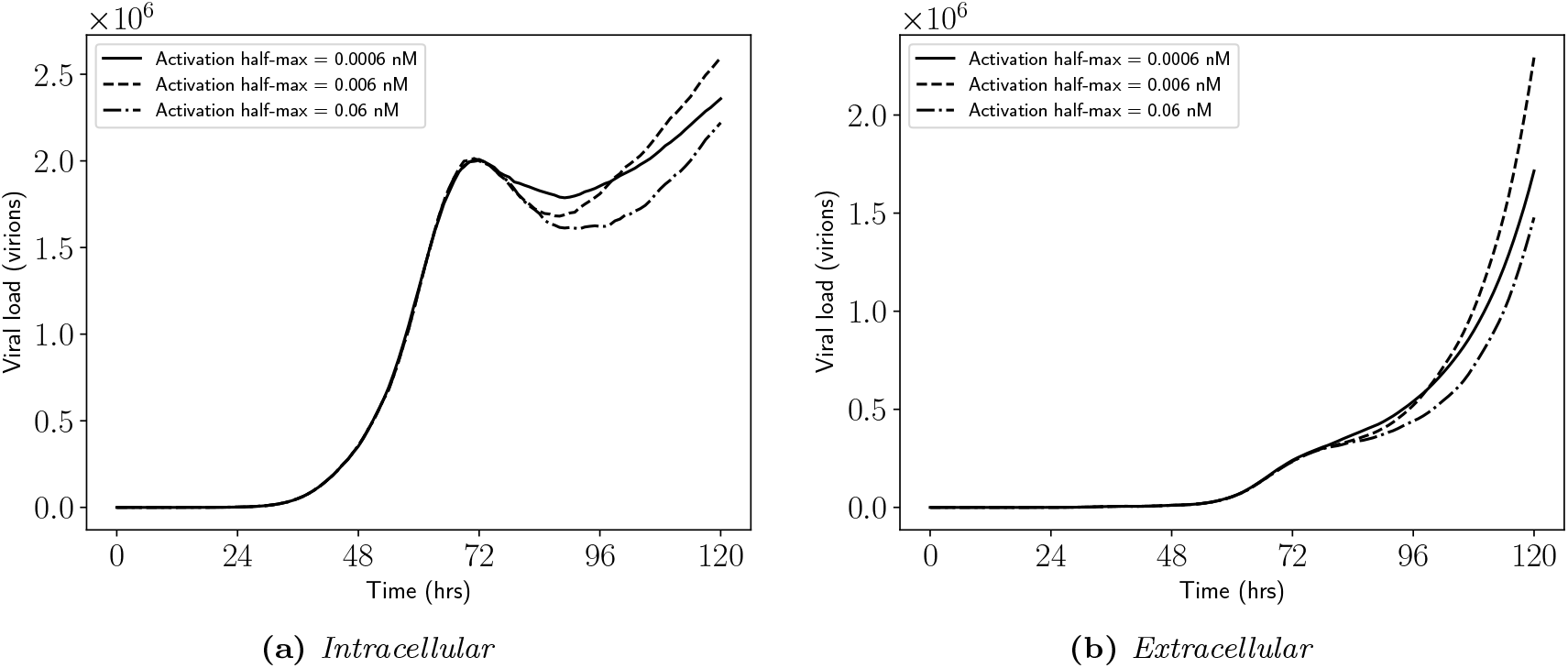
(Viral loads per grid for increasing macrophage activation half-max) *Illustration of (a) intracellular and (b) extracellular viral loads per grid as the simulation progresses, for increasing macrophage activation half-max* 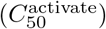. *Note that the MOI* = 0.01 *and the other baseline parameters are given in Tables 1-4*.

Figure 24 illustrates the IFN-I, IL-6, IL-10 and generic mononuclear phagocyte chemokine levels, as the simulation progresses, for each activation half-max value. The trajectories for IFN-I and IL-6 are identical (Figs. 24(a),(b)), where the highest levels of IFN-I and IL-6 are observed for the lowest activation half-max value. The higher levels of IL-10 that are observed for the largest activation half-max value (Fig. 24(c)), are most likely due to the higher numbers of active macrophages observed for the largest activation half-max value. Additionally, higher levels of the generic mononuclear phagocyte chemokine are observed for the largest activation half-max value (Fig. 24(d)), due to the higher total number of apoptotic cells (Fig. 22(b),(f)).

**Figure 24:**
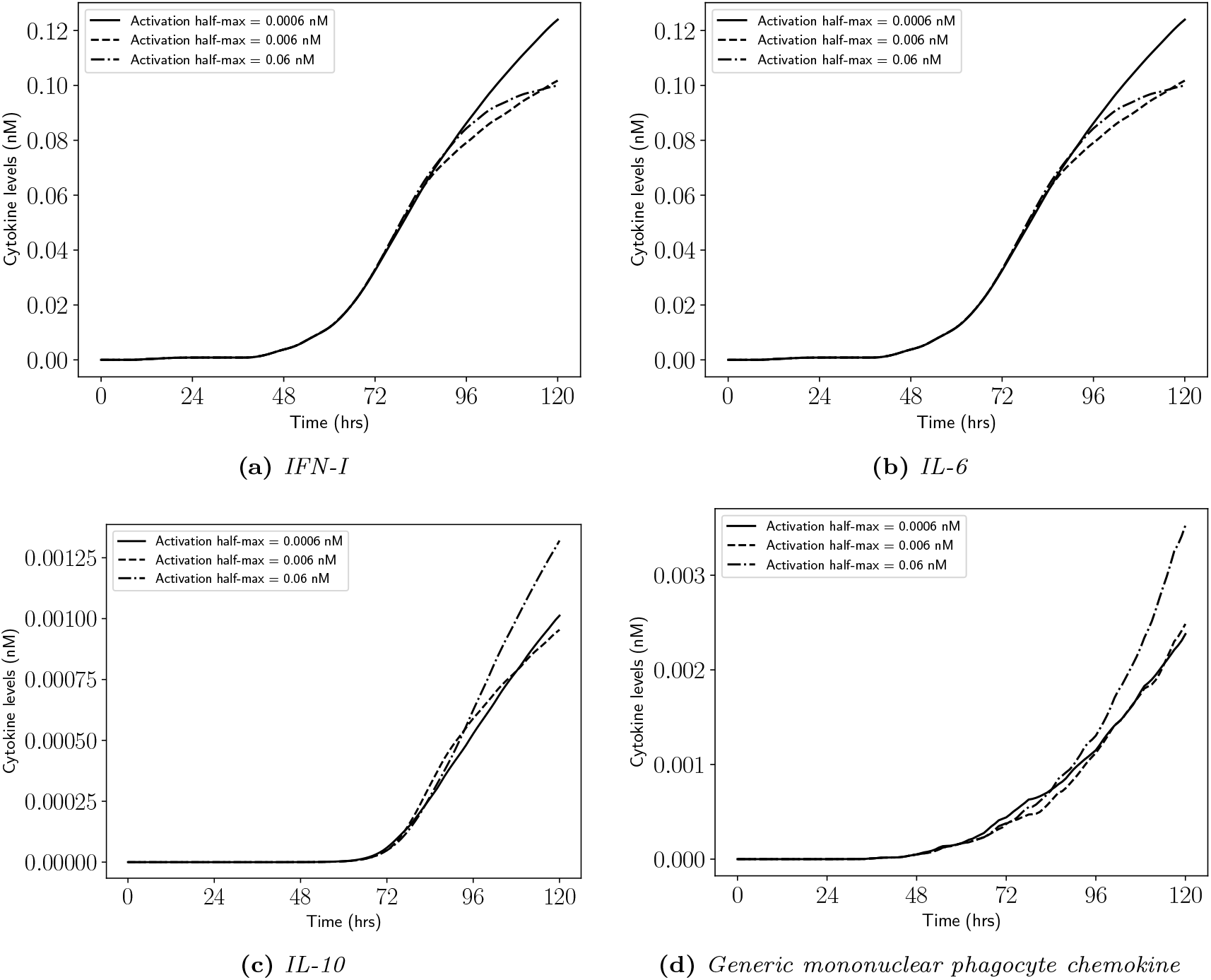
(Cytokine levels for increasing macrophage activation half-max) *Illustration of (a) type I interferon, (b) interleukin 6, (c) interleukin 10 and (d) generic mononuclear phagocyte chemokine levels, as the simulation progresses, for increasing macrophage activation half-max* 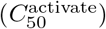. *Note that the MOI* = 0.01 *and the other baseline parameters are given in Tables 1-4*.

## 4 Discussion

The experimental study of host-pathogen dynamics generally involve utilising a combination of cell lines (see e.g. [62]), organoids (see e.g. [63, 64, 65]), animal models (see e.g. [66, 44, 67]), as well as patient and post-mortem data (see e.g. [40, 1, 6, 4]). Mathematical and computational models offer an additional perspective which can supplement experimental studies to enhance the understanding of host-pathogen dynamics (see e.g. [21, 22, 23, 24]). In this article, we have presented a hybrid, multiscale, individual-based model to study the spread of SARS-CoV-2 infection over an epithelial monolayer. We chose to focus on macrophages only, as well as a limited set of cytokines, in order to keep the model complexity to a minimum. The model is then used to study the influence of initial viral deposition, by increasing the initial MOI (Figs. 5-8); the influence of delayed IFN-I secretion from epithelial cells, as well as the magnitude of secretion, by increasing the virus-dependent signal half-max (*S*_50,*V*_) and the secretion rate 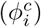 (Figs. 9-16); and the influence of macrophage virus internalisation rate (*r*_[*V*]_), as well as macrophage activation half-max 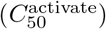 (Figs. 17-24).

Understanding how the initial viral deposition relates to more severe infection and disease outcomes is complicated by the highly nonlinear interactions between the virus, and the innate and adaptive immune systems. Never-the-less, many studies have observed correlations between a high viral load and more severe disease outcomes (see e.g. [1, 68, 69]). Therefore, although our model is limited at this stage (see §4.1), we investigated the role of increasing initial viral deposition (through increasing the initial MOI) on the spread of infection. An increase in infection was observed (Fig. 5), with a higher number of infectious, apoptotic and removed epithelial cells being seen for the highest MOI value considered (Figs. 6(a)-(c)). The increase in infection correlated with an increase in interferon, cytokine and chemokine levels (Fig. 8). A similar correlation has been observed from longitudinal studies (see e.g. [1]). However, care should taken when interpreting the observed correlation; our model is by necessity a simplification, therefore we do not consider other macrophage phenotypes, other immune cell types, or other anti-inflammatory actions, which one would expect to further regulate the interferon, cytokine and chemokine levels. Unexpectedly, we observed a local maximum in the intracellular viral load per grid (Fig. 7), which moved earlier in the simulation as the MOI increased. Within the confines of our model, we hypothesise that a local maximum can only be observed if the export of the intracellular virions exceeds the production (which may happen due to an increase in IFN-I levels relative to the amount of intracellular virus), or if a cluster of infectious epithelial cells have been removed by apoptosis or macrophage phagocytosis, or a combination of both. At the time of the observed local maximum in intracellular viral load per grid, we observed an increase in IFN-I levels but not significant increases in removed or apoptotic epithelial cells, suggesting that the local maximum arises due to IFN-I inhibition of viral entry and replication.

Type I interferons (IFN-I) are potent antiviral cytokines that are produced from a variety cell types in response to a viral infection [42]. Given their potent antiviral action, viruses have evolved to not only evade host-cell detectors so that IFN-I induction is suppressed, but also interfere with interferon-stimulated gene (ISG) induction [14]. Consequently, a delay in IFN-I activity has been suggested to increase disease severity [43, 13], as has been observed for other coronaviruses [44, 45]. Therefore, we investigated the role of increasing delay of IFN-I production from epithelial cells, as well as the magnitude of IFN-I secretion from epithelial cells, on the spread of infection. Increasing the IFN-I secretion delay from epithelial cells increases the spread of infection (Fig. 9), with higher infectious, apoptotic and removed epithelial cells (Figs. 10(a)-(c)) being observed for the longest IFN-I secretion delay. Furthermore, for the longest IFN-I secretion delay, reduced levels of IFN-I were observed (as expected) accompanied by higher levels of IL-6 (Figs. 12(a),(b)). Although increased levels of IL-6 have been consistently implicated in severe COVID-19 (see e.g. [6, 1, 7, 8, 9]), for our model the increase in IL-6 level is merely due to a larger infection. The inclusion of more cytokines and immune cell types, as well as more detailed viral entry and replication pathways, to our model would enable us to investigate the IFN-I and IL-6 relationship more closely, but this is beyond the scope of the current article. Interestingly, for the longest IFN-I secretion delay, we did not observe a local maximum in the intracellular viral load per grid (Fig. 11(a)), further suggesting that IFN-I is responsible for the observed maximum, emphasising its key role in antiviral responses. Moreover, following the local maximum in the intracellular viral load per grid, we observed a rebound from a local minimum (Figs. 7(a), 11(a)), which is commonly observed in low MOI studies and, in our model, is believed to be caused by viral replication inside epithelial cells with low extracellular IFN-I levels. To investigate this further, we studied the role of increasing IFN-I secretion rate on the spread of infection and observed a clear decrease in the spread of infection (Fig. 13). Furthermore, for the highest epithelial IFN-I secretion rate considered, we did not observe a significant rebound in level of intracellular viral load per grid (Fig. 15(a)), enhancing the suggestion that viral replication in the presence of low extracellular IFN-I levels is responsible for the observed rebound in intracellular viral load per grid.

In our model, recruitment of resting macrophages to the site of infection is primarily encouraged by IL-6, with help from the generic mononuclear phagocyte chemokine, and discouraged by IL-10. Therefore, generally, when there is a larger amount of infection, we see higher levels of IL-6 and consequently, greater recruitment of resting macrophages (Figs. 6(d), 10(d), 14(d)). The increase in resting macrophage numbers results in increasing numbers of active and apoptotic macrophages, as expected. To further investigate the role of macrophages on the spread of infection, we varied two macrophage parameters: the macrophage virus internalisation rate, and the macrophage activation half-max. We did not observe increasing infection with increasing macrophage virus internalisation rate (Fig. 17). Furthermore, we did not observe any significant changes in the levels of infectious, apoptotic and removed epithelial cells (Fig. 18); in the levels of intracellular and extracellular viral loads per grid (Fig. 19); or in the cytokine levels (Fig. 20). We argue that this is unsurprising, due to the high viral loads that are observed (e.g. Fig. 19(b)); however, it does potentially suggest that the macrophage virus internalisation rate is either too low or too high to have an observable impact, alternatively it could suggest that the simulation is not sensitive to this parameter, but this cannot be determined without sensitivity analysis, which is beyond the scope of this article. Interestingly, we do observe a difference in resting macrophage recruitment (Fig. 18(d)), even though there are no observable differences in cytokine levels; we attribute this to the stochasticity of the recruitment procedure. On the other hand, for the largest macrophage activation half-max value, we observe a significant influx of resting macrophages. An influx of macrophages/monocytes has been frequently observed in COVID-19 (see e.g. [6, 1, 40, 4]). In our model, we expect the large recruitment of resting macrophages to be a consequence of larger infection and higher IL-6 levels; however, for the largest macrophage activation half-max value, we observe reduced infection (Fig. 22(a)), as well as lower IL-6 levels (Fig. 24(b)), when compared to the smallest macrophage activation half-max value. We do observe a slight increase in generic mononuclear phagocyte chemokine, which could (in combination with IL-6) be enhancing the recruitment of resting macrophages, suggesting a possible feedback between apoptotic cells (epithelial and macrophage) and the recruitment of macrophages, driving tissue injury and inflammation. However, this feedback is likely to be enhanced in our model because resting macrophages do not produce cytokines.

### 4.1 Limitations of this study

The presented model has several limitations: (1) we assume only a single immune cell population, with only a single phenotype; (2) we assume that macrophages may only activate via a single (classical) route; (3) healthy epithelial cells and resting macrophages do not produce cytokines; (4) only a limited set of cytokines is considered; and finally (5) there is a fair amount of uncertainty surrounding the parameter values. Mathematical and computational models require initial simplification, as well as subsequent iterative development, but can never-the-less, offer a unique perspective that can help enhance understanding. Therefore, the presented model serves as an initial starting point from which we can build, by including further complexity and more detailed study which can inform or enhance experimental studies.

## 5 Future Work

When responding to a viral infection, no component of the immune system is expected to work by itself. However, mathematical and computational studies, as well as experimental studies, necessarily require reductions and simplifications, as well as iterative development. Indeed, in this article, we chose to focus only a single type and phenotype of macrophages, as well as a limited set of cytokines and chemokines. Therefore, extending the model to include multiple immune cell populations, as well as subpopulations, and a larger set of cytokines and chemokines is a subject of future work. Furthermore, we chose to focus on a single (classical) activation route, whereas in reality macrophages can alternatively activate; therefore, extending the model to allow for multiple activation routes is also a subject of future work. Multiscale, individual-based models are inherently complex and contain many parameter values which, in general, cannot be expected to directly fit to other modelling work, let alone experimental data. Therefore, improving the parameter estimation and fitting to experimental data, either directly or indirectly via ODE models (see e.g. [70]) is a subject of future work. The presented model has the capability to include more complex intracellular pathways. Therefore, in the future, we would like to study, using more detailed ODE models, the crosstalk between virus replication, IFN-I induction and signalling, as well as pro-inflammatory cytokine induction and signalling, and how the crosstalk impacts the spread of infection. Additionally, the recently discovered Omicron variant seems to display an altered tissue tropism [71, 72], and therefore we would like to consider additional models for virus entry and replication that are specific to different tissue environments. Finally, we would like to take the model to larger scales, either through computational improvements (such as massive parallelisation) or through mathematical models which can *bridge-the-gap* between the scales.

## Footnotes

1 Supplemented by 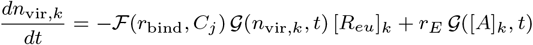

